# Epithelial network percolation reshapes collective mechanochemical dynamics and modes of competitive cell elimination

**DOI:** 10.64898/2026.11.17.767171

**Authors:** Ankita Nayak, Oliver M. Drozdowski, Emmanuel Vikran, Sound WP, Thanh Phuong Nguyen, Edouard Hannezo, Tsuyoshi Hirashima

## Abstract

Living tissues are active materials whose collective states depend on mechanochemical communication across connected cells. Yet tissues undergoing repair, ageing or tumorigenesis often contain intermingled cells with divergent states. How such heterogeneity alters tissue-scale communication, and whether cell competition can restore it, remains unclear. Here we track ERK activity, a dynamic readout and regulator of epithelial mechanochemical coupling, to ask how tissues recover from an aberrant state containing DNA-damaged cells. We show that damaged cells function as insulators, disrupting long-range ERK waves and partitioning control cells into disconnected clusters. As damaged cells are progressively eliminated by neighbouring control cells, fragmented clusters coalesce into a tissue-scale connected network through a percolation transition. Crossing this topological threshold restores collective ERK waves and coordinated cell motion, indicating recovery of tissue-scale mechanochemical order. Percolation also reshapes competitive elimination: before connectivity is restored, control cells remove damaged neighbours through actin-rich invasive protrusions; after percolation, connected control cells eliminate residual damaged cells by ERK-associated multicellular squeezing. Thus, cell competition actively reconfigures tissue topology to restore collective mechanochemical order.

## INTRODUCTION

Biological tissues are self-organized materials in which constituent cells generate forces, remodel adhesions and exchange biochemical signals that feed back to shape tissue-scale structure and dynamics. Through these reciprocal interactions, epithelial sheets acquire collective properties that cannot be inferred from individual cells alone, including coordinated migration, long-range force transmission and tissue-scale signalling patterns that underlie development, homeostasis, wound repair and tissue surveillance^1–4^. Unlike passive materials, living tissues can actively modify the parameters that define their own collective state, including cell density, adhesion, contractility, excitability and network topology^5–8^. A central challenge is therefore to understand how cellular behaviours drive transitions in tissue state, and how these transitions in turn reshape molecular signalling, collective dynamics and cell fate decisions.

Among the cellular processes capable of actively remodelling tissue state, cell competition provides a fundamental mechanism by which tissues detect and eliminate relatively unfit or aberrant cells through local interactions with fitter neighbours. Initially described in developing tissues, this process has been recognized as a broadly conserved tissue-surveillance mechanism that operates in contexts such as oncogenic transformation, differential growth, epithelial repair and DNA damage^9–15^. Recent work has further emphasized the mechanical aspects of cell competition, showing that intercellular force transmission, adhesion strength and dynamic stresses at heterotypic interfaces can influence whether cells survive, die or are extruded^9,16–22^. However, cell competition has largely been considered from the perspective of loser-cell recognition and elimination: which cells are removed, which molecular pathways are involved and how extrusion is executed. Much less is known about the reciprocal question of how the progressive elimination of aberrant cells remodels the physical and topological state of the surrounding tissue, and whether such remodelling alters the tissue’s capacity for collective signalling and self-organization.

A powerful way to probe tissue state as cell collectives is provided by extracellular signal-regulated kinase (ERK) activity^23–28^. In epithelial monolayers, ERK activity forms propagating waves through mechanochemical feedback between cell deformation, ERK signalling, actomyosin contractility and intercellular force transmission ^29–31^. Mechanical stretch can activate ERK, whereas ERK activation promotes contractile responses that deform neighbouring cells, thereby enabling signal propagation across many cell diameters^32–34^. Consequently, ERK waves depend on both mechanical continuity through cell-cell junctions and the ability of cells to convert mechanical inputs into biochemical and contractile responses. ERK dynamics therefore provide not only a biochemical signalling readout, but also a dynamic measure of whether epithelial cells remain sufficiently connected to operate as a coherent mechanochemical collective.

This raises the question of how collective ERK dynamics are altered when epithelial tissues undergo self-organized changes in cellular composition and topology. To address this, we use DNA damage-induced epithelial cell competition as an experimental framework^35^, in which selective elimination of damaged cells progressively transforms a heterogeneous tissue into a more homogeneous state. By mixing control epithelial cells with DNA-damaged cells, we show that damaged cells act as mechanochemical insulators that disrupt intercellular force transmission.

Minimal biophysical modelling predicts a critical dependence of ERK patterning on both the fraction and spatial organization of damaged cells, quantitatively reproducing the transition observed experimentally. We further show that this altered collective state feeds back on competitive cell behaviours: progressive elimination of damaged cells remodels epithelial connectivity until the control-cell population forms a tissue-spanning, percolating network. This transition restores collective ERK dynamics and switches the mode of competitive elimination, linking cell competition, epithelial network connectivity and tissue-scale mechanochemical patterning.

## RESULTS

### DNA-damaged cells suppress emergent ERK activity patterning in epithelial collectives

To introduce DNA damage into epithelial tissues, we treated Madin–Darby canine kidney (MDCK) cells with mitomycin C (MMC), a DNA-crosslinking agent that induces interstrand crosslinks and activates the DNA damage response^36^. Consistent with previous work^35^, transient MMC treatment for 1 hour, followed by washout and reseeding onto culture dishes (Day 0; **Fig. 1A**), induced DNA damage by Day 1. MMC-treated cells exhibited nuclear γ-H2A.X foci and accumulation of phosphorylated p53, whereas dimethyl sulfoxide (DMSO)-treated control cells did not (**Fig. S1A**). Hereafter, we refer to MMC-treated cells after washout as MWO cells, and control DMSO-treated cells as DWO cells.

**Figure 1.**
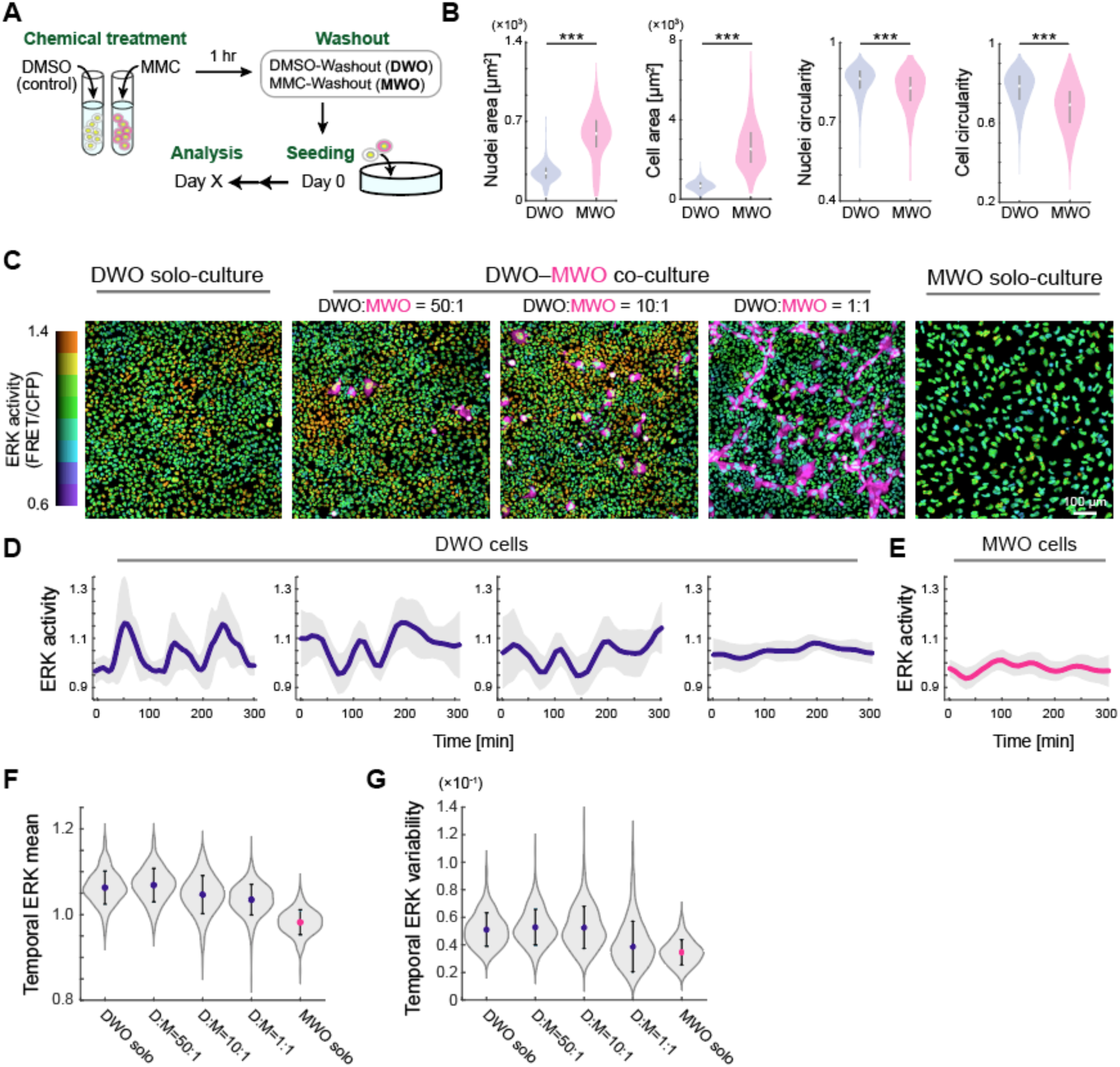
Incorporation of DNA-damaged cells disrupts collective ERK signalling dynamics. (A) Schematic of the experimental workflow. Cells were treated with DMSO as a control or MMC for 1 h, followed by washout to generate DMSO-washout (DWO) and MMC-washout (MWO) cells. The time of reseeding was defined as Day 0, and analyses were performed at the indicated time points. (B) Quantification of nuclear and cellular morphology of DWO and MWO cells on Day 1. Nuclear area, cell area, nuclear circularity, and cell circularity are shown from left to right. n>525 cells per condition from N=3; Welch’s t-test, p<10^-30^ for all cases. (C) Representative snapshots of ERK activity in epithelial monolayers on Day 2 under different culture conditions: DWO solo culture (leftmost), DWO–MWO co-cultures at ratios of 50:1, 10:1, and 1:1 (second to fourth panels), and MWO solo culture (rightmost). The colour bar indicates ERK activity levels. MWO cells are shown in magenta, except in the MWO solo condition, where magenta labelling is omitted for visualization clarity. Scale bar, 100 µm. (D) Temporal dynamics of ERK activity in DWO cells under DWO solo culture and DWO–MWO co-culture conditions (50:1, 10:1, and 1:1). Solid lines represent mean values, and shaded regions indicate the sample standard deviation (s.d). n=84 (DWO solo), n=128 (50:1), n=138 (10:1) and n=286 (1:1). (E) Temporal dynamics of ERK activity in MWO cells under MWO solo culture condition. Solid lines represent mean values, and shaded regions indicate s.d. n=94. (F,G) Quantification of temporal ERK mean (F) and variability (G) in DWO cells across DWO solo and DWO–MWO co-culture conditions, and in MWO cells in MWO solo culture. n>340 cells for each DWO solo and co-culture condition; n=269 cells for MWO solo culture; N=3. See Table S1 for summary of statistical test.

We first characterized the morphological changes of these populations in epithelial monolayers. By Day 1, MWO cells exhibited significantly increased cell and nuclear areas compared to DWO cells (cell: 2.36-fold; nuclei: 3.90-fold) (**Fig. 1B**). In contrast, cell and nuclear circularity were both reduced in MWO cells relative to DWO cells (cell: 0.96-fold; nuclei: 0.88-fold) (**Fig. 1B’**), reflecting increased shape irregularity. We next asked how DNA-damaged cells affect spatiotemporal ERK activity, a dynamic readout and regulator of epithelial tissue state. Using MDCK cells stably expressing a nuclear-localized ERK FRET biosensor^37,38^, we performed live imaging of ERK activity in monolayers established with defined mixtures of DWO and MWO cells (50:1, 10:1, and 1:1) at the time of seeding, Day 0, as well as in corresponding solo-cultures. In DWO monolayers, ERK activity exhibited characteristic propagating waves with spatiotemporal coordination across the tissue, consistent with prior studies of normal epithelial signalling dynamics^29–31^ (**Figs. 1C,D, left; Video 1**). These wave patterns in DWO cells were preserved in co-cultures containing low fractions of MWO cells (50:1 and 10:1), indicating that ERK patterning is robust to sparse perturbations. However, this behaviour collapsed when the fraction of MWO cells increased; in 1:1 co-cultures, DWO cells exhibited spatio-temporally homogeneous ERK activity (**Figs. 1C,D, middle; Video 1**). This homogeneous ERK state was not equivalent to complete ERK inhibition, as pharmacological inhibition further reduced ERK activity in 1:1 co-cultures **(Fig. S1B)**, indicating that the DWO cells maintained an intermediate level of ERK activity despite the loss of propagating ERK waves. MWO cells in solo-culture lacked dynamic ERK patterning, displaying a uniformly low ERK activity profile (**Fig. 1C, right, Fig. 1E; Video 2**).

To quantitatively characterize ERK dynamics at the single-cell level, we calculated the temporal ERK mean, defined as the time-averaged ERK activity of individual cells, and the temporal ERK variability, defined as the standard deviation of ERK activity over time for each cell. In DWO solo cultures and in co-cultures containing a low fraction of MWO cells (50:1), both the temporal ERK mean and temporal ERK variability in DWO cells were comparable to those in DWO solo cultures (**Figs. 1F,G; Table S1** for statistical analyses), confirming that sparse incorporation of MWO cells does not substantially perturb ERK signalling in the tissues. In contrast, increasing the fraction of MWO cells progressively reduced both the temporal ERK mean and temporal ERK variability in DWO cells (**Figs. 1F,G**; **Table S1**). DWO cells in 1:1 co-cultures and MWO cells in solo culture showed significantly lower temporal ERK variability than DWO solo cultures (**Fig. 1G**; **Table S1**), consistent with the loss of clear oscillatory ERK dynamics observed at the tissue level. Together, these results show that increasing proportions of DNA-damaged cells suppress both the average level and temporal fluctuation of ERK activity across epithelial tissues. These findings suggest that coordinated ERK dynamics emerge as a collective property of epithelial cell populations and are disrupted by the incorporation of DNA-damaged cells.

### DNA-damaged cells act as mechanochemical insulators that attenuate intercellular force transmission

To explore how DNA-damaged cells disrupt ERK activity patterning at the tissue level, we first examined cell motility as a functional readout of the mechanochemical coupling. Collective ERK dynamics in epithelial monolayers emerge from coordinated mechanical interactions between neighbouring cells coupled to ERK signalling^32–34,39^. Therefore, we performed single-cell tracking using time-lapse imaging of 1:1 DWO–MWO co-cultures on Day 2 (**Fig. 2A**). Trajectory analysis revealed that DWO cells exhibited persistent displacement over time, whereas MWO cells remained spatially confined with limited net movement (**Fig. 2B**). To quantitatively characterize these differences, we computed the mean squared displacement (MSD), which describes how far cells move through space over time. The temporal scaling of MSD follows a power-law relationship (MSD ∝ t^α^), where the scaling exponent α reflects the persistence of motion (α=1 for diffusive motion; α>1 for super-diffusive, persistent migration). Consistent with the trajectory analysis, DWO cells exhibited a higher scaling exponent (α=1.45), indicative of persistent movement, whereas MWO cells showed a reduced exponent (α=1.32), reflecting more constrained and less persistent dynamics (**Figs. 2C,D**).

**Figure 2.**
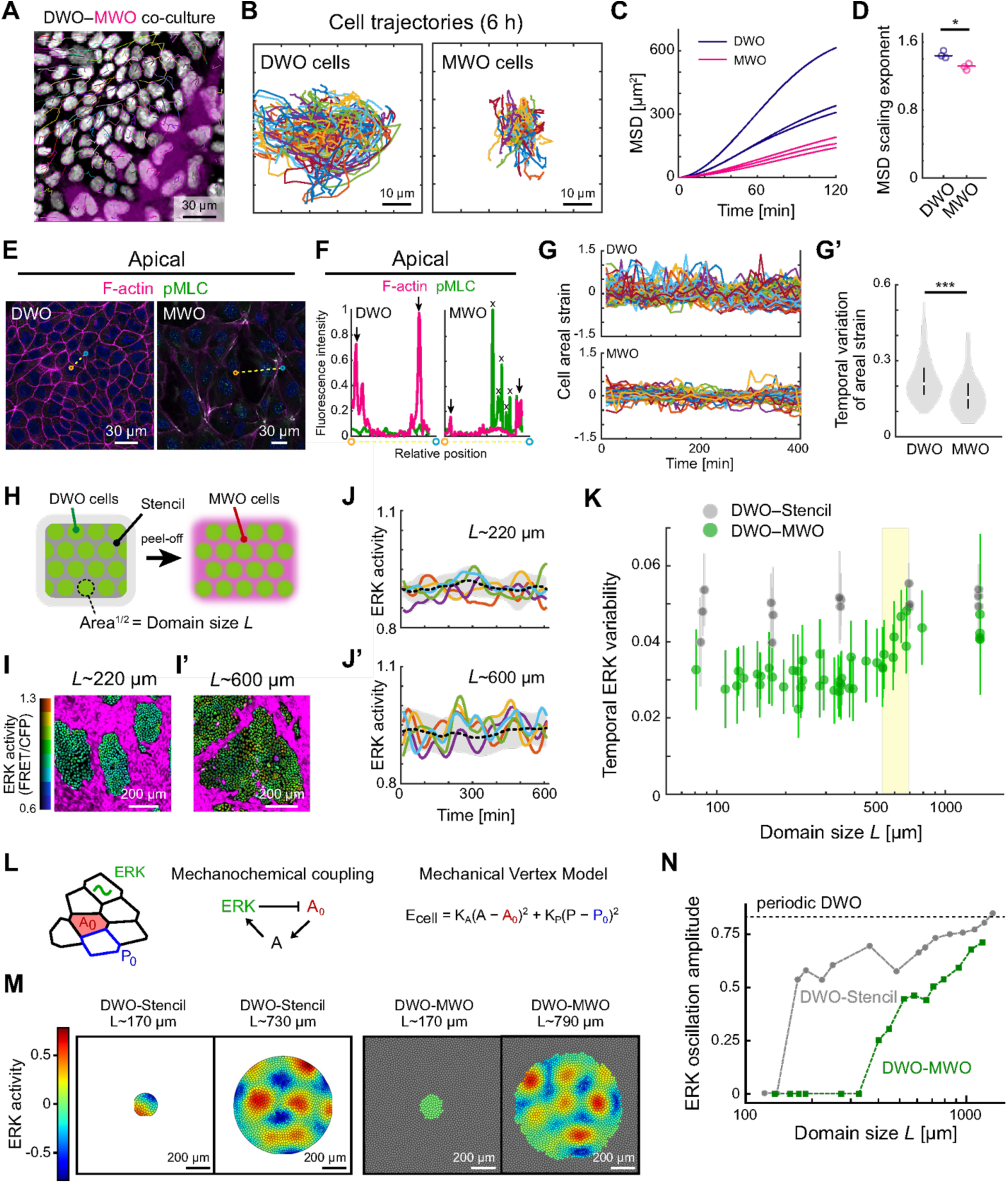
DNA-damaged cells insulate intercellular mechanical communication and impede the emergence of ERK activity patterns. (A) Representative trajectories of DWO and MWO cells over 2 h in 1:1 DWO–MWO co-culture on Day 2. MWO cells are labelled in magenta. Scale bar, 30 µm. (B) Representative trajectories of individual DWO (left) and MWO (right) cells over 6 h on Day 2. Coloured lines represent individual cell trajectories aligned to their initial positions at the center. n>30 cells per condition. Scale bars, 10 µm. (C) Mean squared displacement (MSD) of DWO (blue) and MWO (magenta) cells as a function of time. N=3. (D) MSD scaling exponent for DWO and MWO cells. N=3; Welch’s t-test, p=0.0196. (E,F) Apical distribution of F-actin stained with phalloidin (magenta) and phosphorylated myosin light chain (pMLC; green) in DWO (left) and MWO (right) cells. (E) Fluorescence images. (F) Corresponding intensity profiles along the yellow dotted line indicated in (E). Arrowheads indicate the cell cortex; crosses (x) mark nuclear pMLC puncta in MWO cells. Scale bars, 30 µm. (G,G’) Temporal nuclear areal strain in DWO (top) and MWO (bottom) cells in 1:1 DWO–MWO co-culture. Areal strain was defined as *A*/*A*\*−1, where *A* is the nuclear area and *A*\* is the mean nuclear area for each cell (G). Temporal variability of nuclear areal strain in DWO and MWO cells (G’). n=334 DWO cells and n=173 MWO cells from N=3; two-sample t-test, p=1.51e-13. (H) Schematic of stencil-based micropatterning used to generate spatially confined DWO cell clusters within surrounding MWO cell populations. DWO cells were first seeded into the through-hole regions of the stencil. After stencil removal, MWO cells were seeded into the remaining area of the dish. The cluster domain size *L* was defined as the square root of the projected cluster area. (I,I’) Representative ERK activity maps of confined DWO clusters of different sizes (*L*∼220 µm and *L*∼600 µm) surrounded by MWO cells (magenta). Scale bars, 200 µm. (J,J’) Temporal dynamics of ERK activity in DWO cells confined within clusters of size *L*∼220 µm (J) and *L*∼600 µm (J′). Coloured lines represent individual cells; dashed lines indicate mean values, and shaded regions indicate standard deviation (s.d.). (K) Temporal variability of ERK activity within DWO clusters as a function of domain size *L*. Gray denotes DWO clusters confined by inert stencils (DWO–Stencil), and green denotes DWO clusters surrounded by MWO cells (DWO–MWO). The yellow shaded region highlights the domain size range in which ERK activity variability increases. Error bars indicate s.d. See Table S1 for summary of statistical test. (L) Schematic representation of the mechanochemical vertex model with an internal ERK level which couples mechanically to the target area A_0_ of the vertex model. The cell energy includes an area elasticity term with area stiffness K_A_ and area A, and a perimeter elasticity term with perimeter stiffness K_P_, perimeter P and target perimeter P_0_. (M) Simulated ERK levels in DWO cells confined within small or large circular stencils and in circular DWO domains surrounded by MWO cells. Scale bars, 200 µm. (N) ERK amplitude, defined as half the range between the minimum and maximum ERK level within the domain, plotted as a function of domain size *L*. Grey denotes the DWO-Stencil case, green denotes the DWO-MWO case under periodic boundary condition, and the black dotted line denotes DWO-only simulations under periodic boundary conditions.

We next investigated whether these motility differences were associated with altered cytoskeletal and adhesive organization. Consistent with the morphology evident in **Fig. 1B**, fluorescence staining revealed that MWO cells adopted irregular cell shapes, in contrast to the polygonal organization of DWO epithelial cells. At the apical junctional plane, F-actin and phosphorylated myosin light chain (pMLC) were enriched along DWO cell cortices but were reduced and spatially disorganized in MWO cells (**Figs. 2E,F**). pMLC-positive puncta were also detected in MWO nuclei (**Fig. 2E**), consistent with DNA damage-associated changes in nuclear actomyosin organization^40^. The reduced apical actomyosin organization suggests weakened cortical contractility at MWO cell junctions, which may compromise intercellular force transmission. MWO cells also showed weaker and spatially heterogeneous E-cadherin localization (**Fig. S2A**), consistent with impaired cell–cell adhesion. In line with these observations, temporal area fluctuations were reduced in MWO cells compared with DWO cells (**Figs. 2G,G’**), paralleling the reduced ERK fluctuations observed in MWO cells. Overall, these measurements support that MWO cells exhibit lower mechanochemical activity than DWO cells.

These observations led us to hypothesize that DNA-damaged cells disrupt mechanochemical continuity by acting as insulators within epithelial tissues. To test this, we employed a micropatterning approach to spatially confine DWO cell clusters of defined diameters within surrounding MWO cell populations (**Figs. 2H, S2B**). This system allowed us to systematically vary the bulk-to-boundary ratio and assess its impact on ERK dynamics. In small DWO clusters, ERK activity remained spatially uniform with limited spatiotemporal fluctuations (**Figs. 2I,J; Video 3**). In contrast, larger DWO domains exhibited ERK oscillatory wave patterns with increased amplitude (**Figs. 2I’,J′; Video 4**). Quantitative analysis across domain sizes with one order difference (*L*∼100–1000 µm) revealed that temporal variance of ERK activity increased around 600 µm of the domain size when DWO cells were surrounded by MWO cells, whereas no such size dependence was observed when DWO cells were confined by an inert stencil (**Fig. 2K**; **Table S1**). These results demonstrate that MWO cells attenuate ERK activity amplification in a size-dependent manner, consistent with a boundary-mediated effect. When the DWO bulk region is small relative to the surrounding MWO boundary, ERK patterning is suppressed; conversely, sufficiently large DWO domains overcome this insulating effect and sustain robust ERK dynamics.

To test whether mechanochemical wave disruption is consistent with the observed ERK dynamics, we simulated ERK–mechanical coupling using a vertex model that couples cellular mechanics and ERK signalling^30,41–44^ (**Fig. 2L**). We considered two geometries with different boundary conditions: DWO cells confined by an inert circular stencil, and DWO cell islands surrounded by MWO cells. In the latter case, MWO cells were assumed to be mechanochemical insulators by setting their mechanochemical coupling below the threshold required for oscillations, consistent with our experimental measurements (see Methods). In the stencil condition, ERK waves emerged when the DWO domain was sufficiently large to accommodate the characteristic wavelength (**Fig. 2M**). By contrast, in DWO cell islands surrounded by MWO cells, ERK waves were strongly attenuated, consistent with the experimental observations (**Fig. 2M**; see **Figs. S3A– B’’’** for sensitivity analyses). Consistent with the role of MWO cells as mechanochemical insulators, increasing the effective fluidity of MWO cells reduced wave attenuation. As an additional test of the proposed mechanism, we examined cell dynamics near the boundary in both simulations. Unlike the stencil condition, DWO islands surrounded by MWO cells deformed the adjacent MWO tissue over a short length scale (**Figs. S3C–D’**). This was reflected in movements propagating into the surrounding MWO tissue, whereas stencil-confined DWO domains showed mechanically decoupled tangential flows along the boundary (**Figs. S3E,F**). Particle image velocimetry analysis of experimental time-lapse data revealed flow fields consistent with these predictions (**Figs. S3E’,F’**). Together, these results support a boundary-mediated mechanism in which MWO cells attenuate mechanochemical wave propagation by acting as insulating tissue surrounding DWO domains.

### Elimination of DNA-damaged cells through cell competition establishes percolation of the normal epithelial network

Although MWO cells acted as insulators that disrupted mechanochemical coupling in tissues, DNA-damaged cells are known to be eliminated from epithelial tissues through cell competition^35,45^. Consistent with this, the proportion of MWO cells progressively decreased over time in DWO–MWO co-cultures at both 1:1 and 10:1 mixing ratios, whereas cell composition remained stable in homotypic DWO–DWO cultures (**Figs. 3A–B′, S4A).** MWO cells persisted without substantial loss or detachment in solo culture over comparable durations (**Fig. S4A**), indicating that their loss in the co-culture condition was not due to autonomous cell death or intrinsic instability, but was induced by interactions with neighbouring DWO cells. Immunostaining further showed cleaved caspase-3 in rounded MWO cells after their extrusion from the monolayer (**Fig. S4B**), suggesting that eliminated MWO cells undergo apoptosis and are not reintegrated into the epithelial layer. Together, these observations indicate that MWO cells are selectively eliminated by surrounding DWO cells through cell competition.

**Figure 3.**
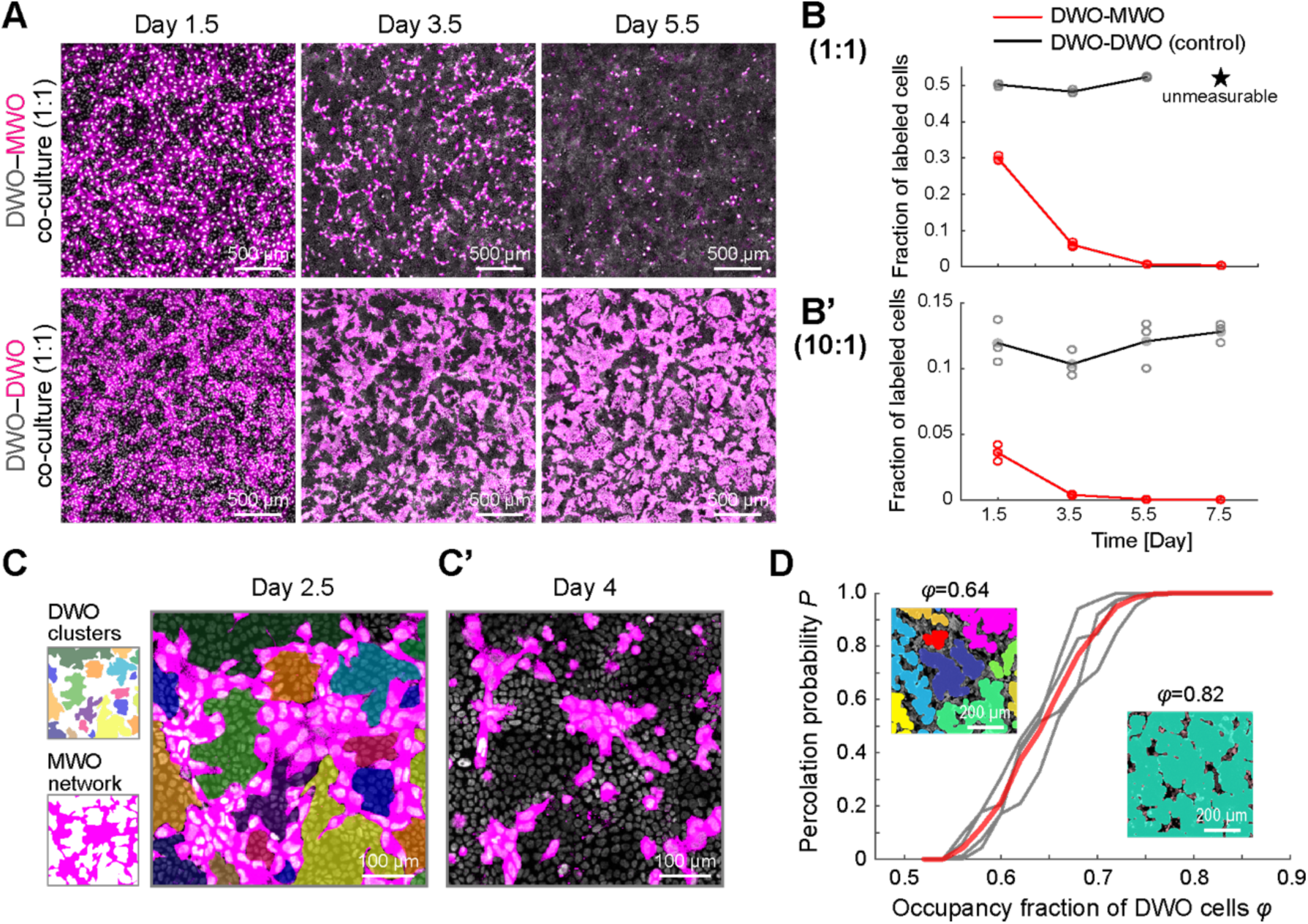
Cell competition establishes percolation of the normal epithelial network. (A) Representative snapshots of the cell competition assay under 1:1 co-culture condition. Top: heterotypic DWO–MWO; bottom: homotypic DWO–DWO (control). Labelled cells are shown in magenta. Images were acquired at Day 1.5, 3.5, and 5.5. Scale bars, 500 µm. (B,B’) Fraction of labelled cells as a function of culture time in 1:1 (B) and 10:1 (B′) co-cultures. Red denotes DWO–MWO, with MWO cells labelled, and black denotes DWO–DWO control, with one of the two cell populations labelled. Each circle represents an independent biological replicate (N=3); solid lines indicate mean values. Asterisks indicate time points where quantification was not possible due to excessive cell density. Jonckheere-Terpstra test, p=6.21e-05 for DWO–MWO and p=0.159 for DWO–DWO (B); p=3.23e-04 for DWO–MWO and p= 0.239 for DWO–DWO (B’). (C,C’) MWO cell network (magenta) and DWO cell clusters on Day 2.5 (C) and Day 4 (C’). In (C), individual DWO clusters are shown in distinct pseudo-colours. Scale bars, 100 µm. (D) Percolation probability (*P*) as a function of the occupancy fraction of DWO cells (*φ*) in 1:1 DWO–MWO co-culture. Individual measurements from four samples (N=4) are shown in gray, calculated over the time window Day 2.5–4.0; the red line indicates the mean. Insets show representative images of connected DWO clusters, with colours indicating distinct clusters, at *φ*=0.64 (left) and *φ*=0.82 (right). Scale bars, 200 µm.

This progressive elimination was accompanied by large-scale reorganization of tissue topology. At early time points, such as Day 2.5, MWO cells formed a contiguous network that partitioned the epithelial sheet into multiple isolated DWO clusters (**Fig. 3C**). Over time, this MWO network progressively collapsed, allowing neighbouring DWO clusters to merge. By Day 4, this process generated a giant DWO cluster that spanned the field of view from edge to edge (**Fig. 3C′**). This behaviour is reminiscent of a percolation transition, in which increasing local connectivity leads to the emergence of a system-spanning cluster^46,47^. Motivated by this, we quantified the occupancy fraction of DWO cells, *φ*, over time from timelapse imaging data. As expected from the progressive elimination of MWO cells, *φ* increased monotonically over time (**Fig. S4C**). We then computed the percolation probability *P* as a function of *φ* to assess global DWO-cell connectivity. The resulting percolation probability curve exhibited a sharp sigmoidal transition (**Fig. 3D**), indicating a rapid shift from fragmented DWO clusters to a connected DWO network. The probability of forming a spanning DWO cluster reached 95% at *φ*=0.72, defining a critical threshold for percolation in this system. Because *φ* increases over time, this threshold also allowed us to define the timing of the percolation transition in each dataset. We therefore normalized time relative to this transition point, hereafter referred to as the percolation time, to compare cellular and tissue dynamics before and after percolation. This framework separated the co-culture process into pre-percolation and post-percolation regimes, enabling us to examine how normal-cell connectivity regulates ERK dynamics and the mode of DNA-damaged cell elimination.

### Cell competition-induced percolation triggers a transition in ERK activity patterns and cell dynamics

We then examined how ERK activity evolves across this transition by long-term live imaging of ERK activity initiated at Day 2.5 in DWO–MWO co-cultures. During the pre-percolation phase, ERK activity in DWO cells remained relatively uniform, consistent with the previous result (**Fig. 1C**); however, upon crossing the critical percolation time, ERK activity began to exhibit amplified fluctuations, giving rise to coherent oscillatory wave patterns (**Fig. 4A; Video 6**). The ERK oscillatory period, approximately 100 min (**Figs. 4B,B’**), was comparable to that observed in intact epithelial monolayers reported previously^30,31^, indicating that the intrinsic ERK oscillatory machinery is preserved but becomes dynamically engaged only after the percolation transition. Consistent with this, conditions with a low fraction of MWO cells, where insulating boundaries are reduced, also supported robust ERK pattern formation. These results suggest that the emergence of ERK activity patterns is governed by connectivity within the tissue. Specifically, the breakdown of the MWO cell network eliminates insulating boundaries, enabling long-range mechanical coupling among DWO cells. This restoration of connectivity allows mechanochemical feedback to propagate across the tissue, thereby amplifying and sustaining ERK oscillations.

**Figure 4.**
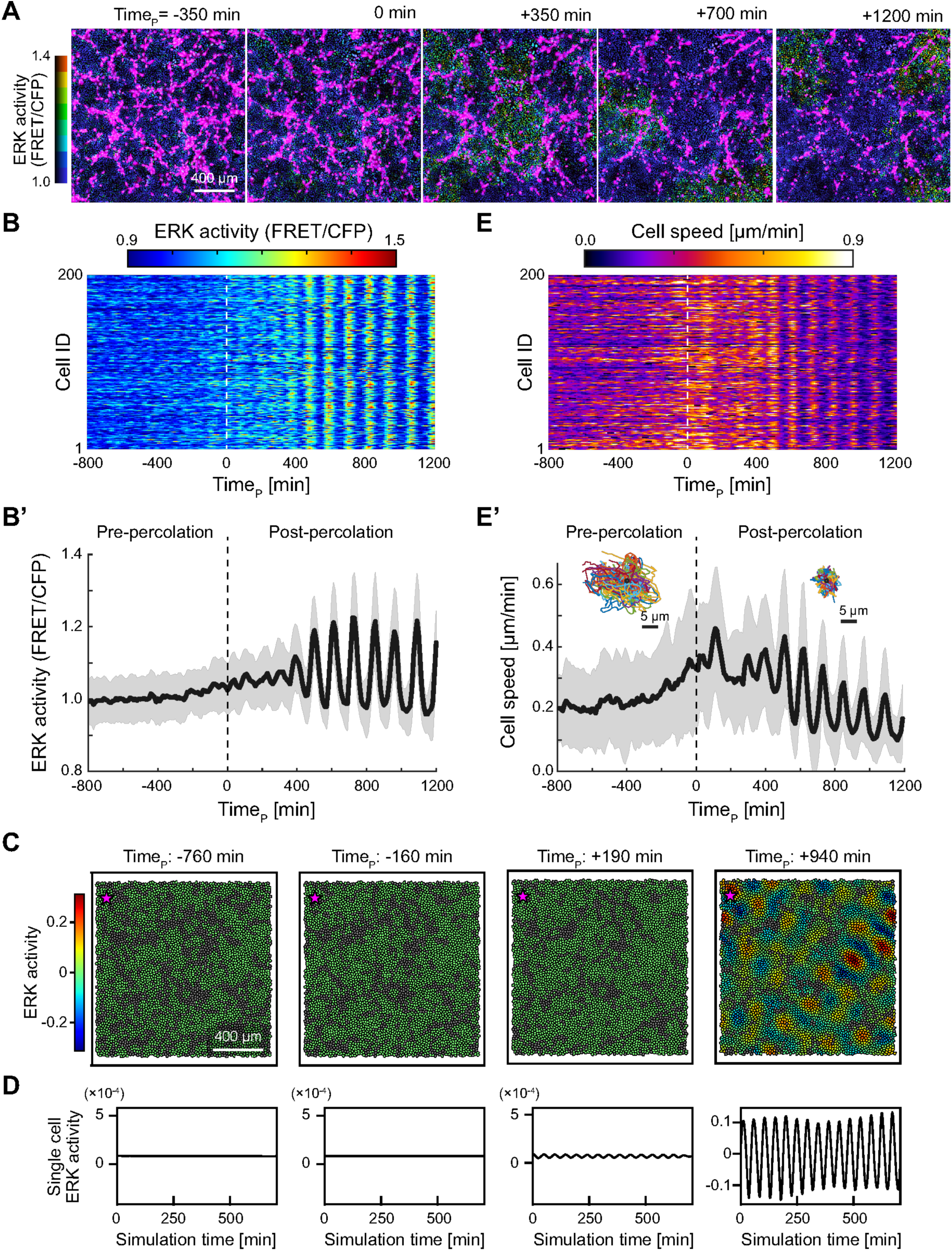
Cell competition-induced percolation drives ERK activity patterning. (A) Snapshots of time-lapse images of ERK activity map in 1:1 DWO–MWO co-culture. Time stamp on the top represents the critical percolation time. Scale bars, 400 µm. (B,B’) Heatmap of ERK activity in 200 individual cells sampled from a representative local region (B); rows correspond to cell IDs and columns to normalized time. The corresponding median ERK activity is shown as a thick line, with the standard deviation (s.d.) indicated by the shaded area (B′). (C) Simulated ERK activity in DWO–MWO cellular arrangements extracted from cropped experimental fields at the indicated start times relative to the critical percolation time. Cell positions and identities were kept fixed during each simulation. Scale bars, 400 µm. (D) Simulated single-cell ERK activity traces for the representative cell marked by a magenta star in (C). (E,E’) Heatmap of instantaneous cell speed in the same 200 cells as in B (E); rows correspond to cell IDs and columns to normalized time. The corresponding median speed is shown as a thick line, with the s.d. indicated by the shaded area (E′). Insets show representative cell trajectories during the pre-percolation (left) and post-percolation (right) phases, after aligning the initial position of each trajectory to a common origin (circle). Scale bars, 5 µm.

To test whether the onset of ERK waves could be explained by the changing topology of the DWO–MWO network, we performed data-driven simulations using cellular arrangements extracted from experimental time-lapse images. In these simulations, MWO cells were treated as insulating cells and distributed according to MWO masks at selected time points before and after the critical percolation time, while cell positions and identities were kept fixed during each simulation. Using the same model parameters as above, ERK dynamics remained suppressed in pre-percolation arrangements, where DWO cells were fragmented by a dense MWO network (**Figs. 4C,D; Video 7**). By contrast, post-percolation arrangements supported self-sustained ERK oscillations and propagating wave-like patterns, consistent with the experimentally observed onset of ERK waves after DWO-domain percolation. Thus, the emergence of ERK waves can be reproduced by the temporal reorganization of network connectivity, supporting that cell competition-induced percolation triggers coordinated changes in ERK activity and cellular dynamics.

Not only the ERK activity but also cellular dynamics are similarly regulated by the percolation transition. Prior to percolation, cell motility progressively increased, with cells exhibiting more persistent and coordinated movement as the system approached the critical point (**Figs. 4E,E′**). Following percolation, instantaneous cell speed displayed oscillatory behaviour that was temporally coupled to ERK activity, consistent with known coupling between ERK signalling and contractile force generation^27,31,34^. Notably, over longer timescales, cell motility gradually decreased and cell position became more spatially confined, indicating a transition toward a mechanically constrained state. This behaviour suggests a shift in the collective mechanical regime of the tissue, consistent with a rigidity transition from a dynamically fluctuating state to a more restricted, solid-like state^48,49^. Collectively, these findings demonstrate that cell competition drives a percolation transition of DWO cell connectivity, which in turn triggers coordinated changes in ERK activity and cellular dynamics.

### Normal-cell protrusions mediate DNA-damaged cell elimination during the pre-percolation

We next asked how DWO cells eliminate MWO cells during the pre-percolation phase. To visualize cell morphology, we used MDCK cells expressing Lifeact–mCherry to label F-actin in either DWO or MWO cells (F-actin^DWO^ or F-actin^MWO^) under co-culture conditions. In homotypic DWO cultures (i.e., DWO–DWO co-culture), cell-cell interfaces displayed the typical polygonal junctional organization characteristic of epithelial monolayers as anticipated (**Fig. 5A, left**). In heterotypic co-cultures (i.e., DWO–MWO co-culture), DWO cells in contact with MWO cells extended elongated F-actin–rich protrusions toward adjacent MWO cells (**Fig. 5A, middle**). MWO cells in contact with DWO cells also formed similar protrusions toward DWO cells (**Fig. 5A, right**), indicating that both cell types respond to heterotypic interfaces by generating protrusive structures. Quantification showed that the number of protrusions per interface-adjacent cell was increased in DWO–MWO co-cultures compared with homotypic DWO cultures (**Fig. 5B**). Because these actin-based protrusions were enriched at heterotypic interfaces, we refer to them as EpiArms (elongated protrusions induced through actin dynamics for recognizing mismatched cells).

**Figure 5.**
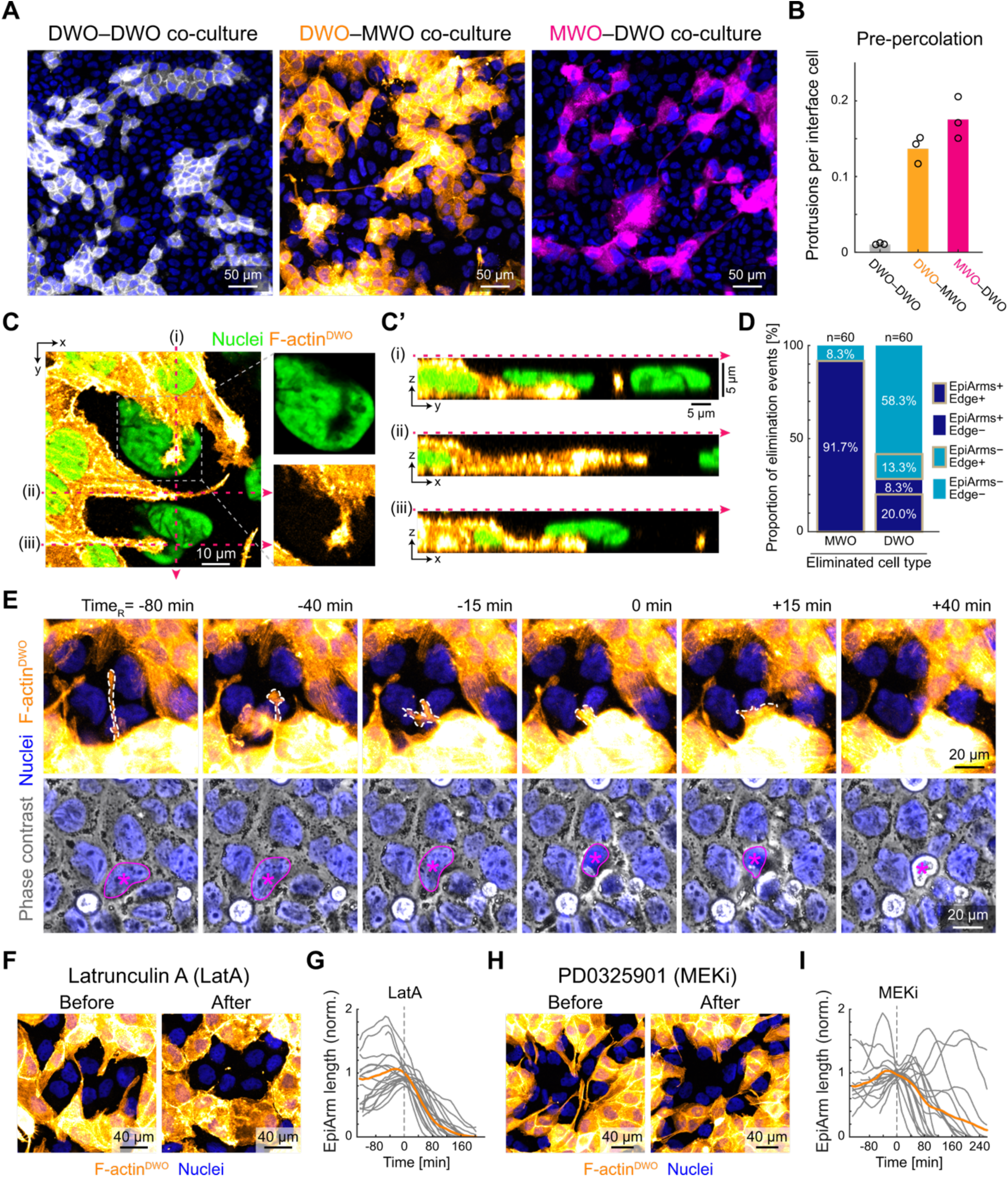
MWO cells are eliminated by EpiArms during pre-percolation phase. (A) Representative images of homotypic DWO–DWO co-culture (left) and heterotypic DWO–MWO (middle) and MWO–DWO (right) co-cultures with nuclei counterstained on Day 2.5. DWO cells are shown in white (left) or orange (middle), and MWO cells in magenta (right). Scale bars, 50 µm. (B) Protrusion frequency per labelled cell at the interface with unlabelled cells. Bars represent mean values. N=3. (C,C′) Representative images illustrating physical interactions between DWO EpiArms and MWO cells. DWO cells are labelled with Lifeact–mCherry (orange; F-actin^DWO^), and MWO cells are unlabelled with nuclei counterstained in green. The boxed region (dashed line) is magnified in (C′), showing the MWO nucleus (top) and the DWO EpiArm (bottom). (C) shows the xy plane, with corresponding orthogonal views along axes (i–iii) indicating xz or yz sections in (C’). Scale bars, 10 µm (C) and 5 µm (C′). (D) Quantification of EpiArm-associated elimination events for MWO cell removal events with F-actin^DWO^ (left) and for DWO cell removal events with F-actin^MWO^ (right). Stacked bar plots showing the proportions of elimination events classified into four categories based on the presence or absence of EpiArms and localization at heterotypic edges: EpiArm+/Edge+, EpiArm+/Edge−, EpiArm−/Edge+, and EpiArm−/Edge−. n = 60 elimination events for each condition from N=3; chi-square test for the full 2×4 contingency table, p= 1.28e-13. (E) Time-lapse imaging of MWO cell elimination driven by actin-based protrusions (EpiArms) from DWO cells. Upper panels show DWO cells labelled with Lifeact–mCherry (orange; F-actin^DWO^) and unlabelled MWO cells with nuclei counterstained in blue. Dotted lines indicate the EpiArm involved in MWO cell elimination. Lower panels show corresponding phase-contrast images; asterisks mark the position of the MWO nucleus, and outlines indicate nuclear deformation during extrusion. Time 0 min is defined as the onset of nuclear shrinkage preceding extrusion. Scale bars, 20 µm. (F) Morphological changes in DWO EpiArms (orange) before and after treatment with 100 nM Latrunculin A (LatA). MWO cells are unlabelled, with nuclei counterstained in blue. Scale bars, 40 µm. (G) Time course of EpiArm length normalized to its value immediately before LatA treatment. Gray lines represent individual measurements, and the orange line indicates the mean. Time 0 denotes the time of treatment. n=19, N=3. (H) Morphological changes in DWO EpiArms (orange) before and after treatment with 1 µM PD0325901 (MEK inhibitor, MEKi). MWO cells are unlabelled, with nuclei counterstained in blue. Scale bars, 40 µm. (I) Time course of normalized EpiArm length following MEKi treatment. Gray lines represent individual measurements, and the orange line indicates the mean. Time 0 denotes the time of treatment. n=18, N=3.

To determine how EpiArms interact with neighbouring cells within the epithelial layer, we analyzed their three-dimensional organization. EpiArms extended beneath adjacent MWO cells, penetrating the basal region and locally deforming the nuclei of target cells (**Figs. 5C,C′**). Consistent with this, in confocal sections through the nuclear plane, regions occupied by EpiArms showed strong F-actin signal with a corresponding exclusion of nuclear signal, indicating that EpiArms extend beneath the target cell body and locally indent the nucleus (**Fig. 5C, inset**). During extension, some EpiArms partially detached from the substrate and traversed through lateral intercellular spaces (**Fig. 5C’ii**), indicating that they can navigate confined regions within the epithelial layer while exerting forces capable of deforming neighbouring cells.

We then examined whether EpiArms were associated with cell elimination by classifying removal events according to EpiArm presence. During MWO cell elimination, 91.7% of events were accompanied by DWO-derived EpiArms at heterotypic interfaces (**Fig. 5D, left**). The remaining EpiArm-negative MWO elimination events occurred away from heterotypic interfaces. By contrast, during DWO cell elimination, 71.6% of events occurred without EpiArms (**Fig. 5D, right**). The distribution of event categories differed significantly between MWO and DWO elimination, with EpiArm-positive events enriched during MWO removal. These results suggest that DWO-derived EpiArms preferentially contribute to MWO cell elimination, whereas DWO cell elimination occurs largely independently of MWO-derived EpiArms. Time-lapse imaging further revealed that EpiArms undergo repeated cycles of extension and retraction, during which they infiltrate the basal region of neighbouring MWO cells (**Video 8**). Notably, nuclear shrinkage in MWO cells occurred preferentially during the retraction phase of EpiArms (**Fig. 5E**). As nuclear shrinkage is an early marker associated with cell elimination^24^, this temporal relationship suggests that EpiArm retraction contributes to the initiation of MWO cell removal.

To test the requirement of EpiArms, we disrupted actin polymerization using Latrunculin A (LatA). LatA treatment led to rapid retraction and loss of EpiArms within 2 h (**Figs. 5F,G,S5A; Video 9**). Further, long-term heterotypic co-culture assays revealed that LatA treatment significantly reduced MWO cell elimination (**Fig. S5B**), supporting a requirement for actin-dependent protrusive activity in efficient removal of DNA-damaged cells. We also examined whether EpiArm formation depends on ERK activity. Inhibition of ERK signalling using a MEK inhibitor (MEKi, PD0325901) suppressed both the extension and maintenance of EpiArms (**Figs. 5H,I,S5C; Video 10**). Given that ERK activity remains at an intermediate level during the pre-percolation phase in DWO–MWO co-cultures (**Figs. 1C,D,S1B**), these results suggest that intermediate ERK activity may support, rather than simply accompany, sustained EpiArm dynamics.

### Normal-cell percolation switches the mode of DNA-damaged cell elimination

Although EpiArms were enriched during the pre-percolation phase, they were rarely observed after DWO cells formed a percolating network. In the post-percolation phase, DWO cells surrounding small clusters of MWO cells formed smooth heterotypic interfaces, without the elongated protrusions seen before percolation (**Fig. 6A**). We then examined the organization of actomyosin at these heterotypic cell interfaces in the post-percolation phase. F-actin was enriched at the apical DWO–MWO interface (**Fig. 6B, left**), whereas the active myosin marker pMLC accumulated in DWO cells near the interface, particularly at the basal plane (**Fig. 6B, right**). This organization is consistent with increased contractile tension along the heterotypic boundary. In addition, basal pMLC accumulation beneath the nucleus, which was prominent in MWO cells during the pre-percolation phase (**Figs. S6A,B**), was absent after percolation (**Fig. 6B, right**), suggesting reduced engagement between MWO cells and substrate.

**Figure 6.**
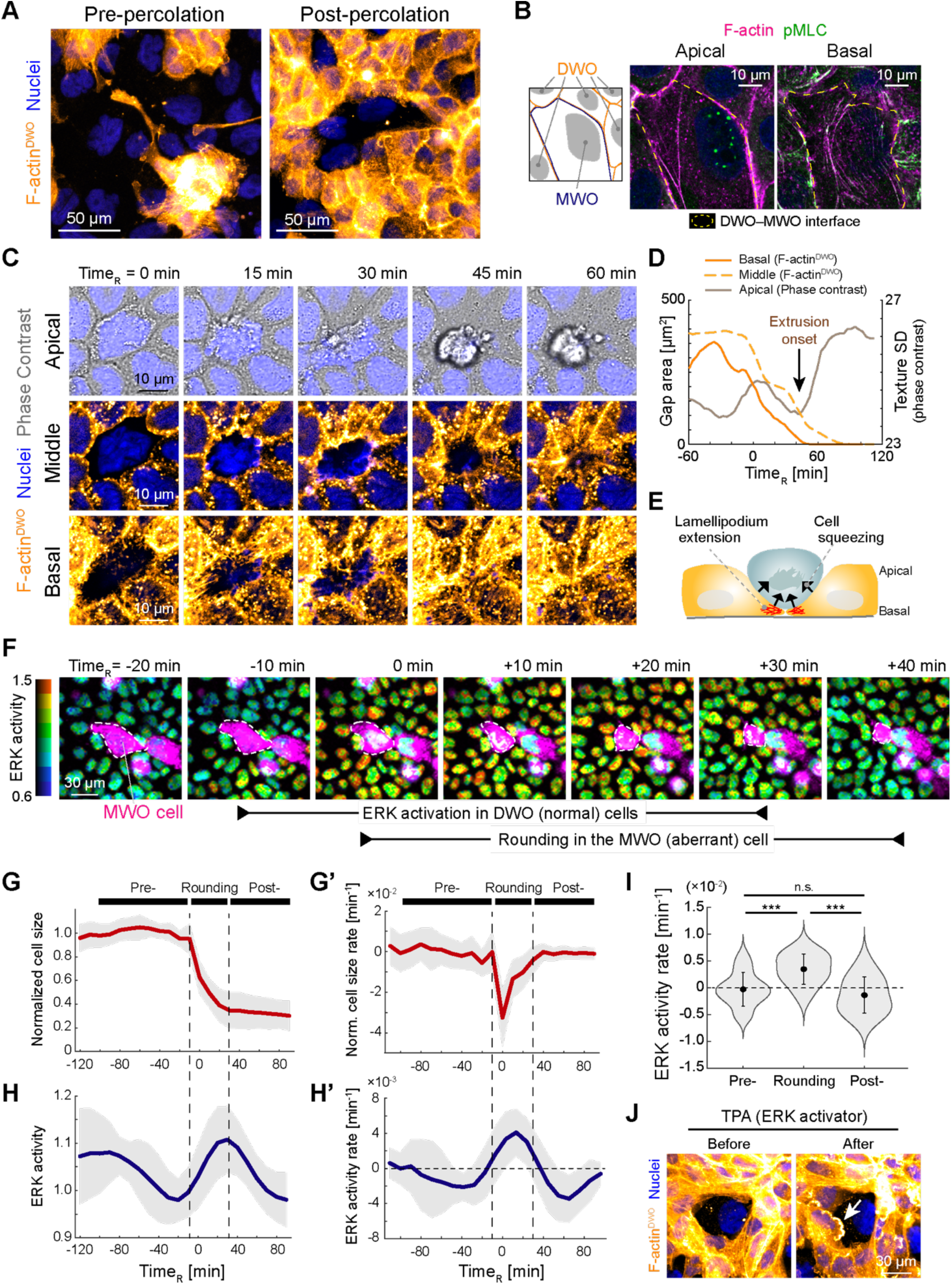
Percolation of normal cells shifts the mode of cell competition. (A) Representative images of DWO–MWO co-cultures during the pre-percolation (left) and post-percolation (right) phases. DWO cells are labelled with Lifeact–mCherry (orange; F-actin^DWO^), whereas MWO cells are unlabelled. Nuclei are counterstained with CFP (blue). Scale bars, 50 µm. (B) Distribution of F-actin stained with phalloidin (magenta) and phosphorylated myosin light chain (pMLC; green) at apical and basal planes in a DWO–MWO co-culture at Day 4. As illustrated schematically (leftmost), a single MWO cell is surrounded by DWO cells. Yellow dashed lines indicate the DWO–MWO interface. Scale bars, 10 µm. (C) Time-lapse imaging of MWO cell elimination driven by squeezing from surrounding cells during the post-percolation phase. DWO cells labelled with Lifeact-mCherry (orange; F-actin^DWO^) surrounding an unlabelled MWO cell in the center. Nuclei are counterstained with CFP (blue). Top panels show phase-contrast images at the apical plane; middle and bottom panels show F-actin signals in DWO cells at middle and basal planes, respectively. Time 0 is defined as the onset of MWO nuclear shrinkage. Scale bars, 10 µm. (D) Temporal changes in gap area at the basal and middle planes (left axis) and texture standard deviation (SD) of phase-contrast images at the apical plane (right axis). Time 0 is defined as the onset of nuclear shrinkage. The onset of a progressive increase in texture SD corresponds to the onset of cell extrusion. (E) Schematic of MWO cell squeezing and extrusion driven by lamellipodial extension of surrounding DWO cells. (F) Time-lapse snapshots showing elimination of an MWO cell (magenta) and ERK activity in surrounding DWO cells (colour bar shown on the left) during the post-percolation phase. The dashed outline marks the MWO cell undergoing elimination. Time 0 is defined as the onset of MWO cell shrinkage. Scale bar, 30 µm. (G,G′) Temporal changes in MWO cell size normalized to its value at the onset of rounding (G) and the corresponding temporal rate (G′). Red lines indicate mean values, and shaded regions indicate standard deviation (s.d.). n=20, N=4. (H,H′) Temporal changes in ERK activity in surrounding DWO cells normalized to the value at the onset of MWO cell rounding (H) and the corresponding temporal rate (H′). Blue lines indicate mean values, and shaded regions indicate s.d. n=20, N=4. (I) Temporal rate of ERK activity across pre-rounding, rounding, and post-rounding phases. n=60, N=4 for each condition. See Table S1 for statistical details. (J) Snapshots of F-actin organization before and after treatment with 10 nM TPA during post-percolation phase. DWO cells are labelled with Lifeact–mCherry (orange; F-actin^DWO^), whereas MWO cells are unlabelled. Nuclei are counterstained with CFP (blue). The arrow indicates lamellipodial extension. Scale bar, 30 µm.

Because MWO cells continued to be eliminated after percolation despite the loss of EpiArms (**Figs. 3B,S4C,6A**), we next analyzed the extrusion process in detail. We co-cultured Lifeact–mCherry– expressing DWO cells with unlabelled MWO cells and performed live-cell imaging to monitor F-actin dynamics (**Video 11**). During the post-percolation phase, DWO cells surrounding an MWO cell extended basal lamellipodia beneath the target cell, and these DWO lamellipodia progressively covered the basal adhesive area of the MWO cell (**Fig. 6C**). Following lamellipodial extension, the F-actin–enriched interface at the mid-apical plane gradually constricted, leading to closure of the surrounding DWO cell ring and extrusion of the MWO cell (**Fig. 6C**). Quantification showed that basal gap closure preceded closure at the middle plane and nuclear shrinkage in MWO cells; both basal and middle-plane closure showed a progressive reduction in area during extrusion, and MWO cell extrusion began after complete closure of the basal gap (**Fig. 6D**). Thus, post-percolation elimination occurs through a coordinated squeezing mechanism initiated by basal lamellipodial extension (**Fig. 6E**), rather than through an EpiArm-mediated regime.

We then asked what initiates the basal lamellipodial extension and contractile squeezing that drive MWO cell extrusion after percolation. Because ERK activity waves emerged during the post-percolation phase and ERK signalling is linked to force generation, we examined whether ERK activity in surrounding DWO cells was temporally coupled to MWO cell extrusion. We aligned ERK activity traces to the onset of MWO cell rounding, which marked the beginning of extrusion, and quantified ERK dynamics in surrounding DWO cells over a time window spanning before and after this event (**Fig. 6F; Video 12**). ERK activity in DWO cells increased before MWO cells began to round and remained elevated as MWO cells progressed toward extrusion from the epithelial monolayer. During rounding, the projected area of MWO cells progressively decreased to 30.2% of the initial area (**Fig. 6G**). To define the extrusion sequence quantitatively, we calculated the temporal derivative of MWO cell area. This analysis separated the process into three phases: pre-rounding, rounding and post-rounding (**Fig. 6G’**). The transition from pre-rounding to rounding was defined by the onset of sustained negative area change. We then compared ERK dynamics in surrounding DWO cells across these phases. ERK activity began to increase during the pre-rounding phase, before the onset of MWO cell rounding. On average, the temporal derivative of ERK activity crossed zero approximately 10 min before the transition from pre-rounding to rounding (**Figs. 6H,H’**). Consistently, the mean rate of ERK activity change was positive during the rounding phase, whereas it was close to zero or negative during the pre-rounding and post-rounding phases (**Fig. 6I**). These results indicate that ERK activation in surrounding DWO cells precedes and accompanies MWO cell rounding during post-percolation extrusion.

To test whether acute ERK activation is sufficient to induce heterotypic interface remodelling, we stimulated ERK signalling in DWO–MWO co-cultures using either the small-molecule activator 12-O-tetradecanoylphorbol-13-acetate (TPA) or recombinant epidermal growth factor (EGF). Treatment with either activator increased lamellipodial extension in DWO cells surrounding MWO cells at heterotypic interfaces (**Figs. 6J,S6C**). This response was observed in the post-percolation phase and also in the pre-percolation phase, where EpiArms were normally enriched. Under pre-percolation conditions, acute ERK activation induced plasma membrane ruffling and lamellipodium-like basal extensions but did not suppress EpiArm formation at DWO–MWO interfaces (**Figs. S6D,D’; Videos 13,14**). These results indicate that acute ERK activation promotes basal plasma membrane extensions, but is not sufficient to explain the disappearance of EpiArms after percolation. Together, these data support that ERK activation in DWO cells preceding MWO cell rounding promotes basal lamellipodial extension at heterotypic interfaces. Thus, after DWO cells form a percolating network, ERK activity dynamics are associated with a switch in the dominant mode of MWO cell elimination, from EpiArm-mediated basal invasion to lamellipodial coverage followed by contractile squeezing.

## DISCUSSION

In this study, we used a minimal epithelial co-culture system to ask how competitive elimination of aberrant cells reshapes tissue state over time. Mixing control (DWO) and DNA-damaged (MWO) epithelial cells generated a heterogeneous monolayer in which damaged cells initially disrupted mechanochemical coordination, yet were progressively eliminated by surrounding control cells. Because this simple in vitro setting preserved epithelial connectivity and ERK-based mechanochemical signalling while limiting cellular complexity, it allowed us to follow ERK activity, cell motion, network topology and elimination behaviours continuously across the transition from a mixed population to a connected normal-cell network. We found that cell competition does more than remove damaged cells: it remodels epithelial connectivity through a percolation transition, restoring collective mechanochemical dynamics and changing the mode of cell elimination. This transition differs from previously described jamming, rigidification and fluid-to-solid transitions, in which the material state of the tissue as a whole changes through alterations in cell density, shape, motility or adhesion^48,50–55^. Here, the epithelial monolayer remains confluent, whereas the connectivity of the normal-cell subnetwork is progressively reorganized. As MWO cells are eliminated, initially fragmented DWO clusters coalesce into a tissue-spanning network, producing a cell-type-specific percolation transition. This network is neither imposed externally nor generated by passive tissue growth, but actively constructed through competitive elimination. Cell competition can therefore act as a topological remodelling process, restoring long-range mechanochemical communication through the selective removal of mechanically disruptive cells.

This framework also changes how cell competition itself can be viewed. Our results suggest an additional tissue-level consequence: as loser cells are progressively removed, the surrounding winner tissue crosses a connectivity threshold that alters the conditions under which further elimination occurs. Before percolation, DWO cells exist as isolated or weakly connected clusters separated by an MWO network. Under this topology, DWO cells cannot consistently surround individual MWO cells or MWO clusters, making collective squeezing inefficient. Instead, MWO removal occurs predominantly through EpiArms, elongated actin-rich protrusions that extend from DWO cells into neighbouring MWO cells. Thus, pre-percolation elimination can proceed through local one-to-one interactions at heterotypic interfaces, without requiring complete encirclement of the target MWO cell by DWO neighbours. After percolation, by contrast, DWO cells form a connected network that surrounds residual MWO cells or clusters, creating a geometry in which basal lamellipodial coverage and collective contractile squeezing become effective. Thus, the mode of competitive elimination is not fixed, but emerges from the connectivity and geometry of the tissue state.

EpiArms represent a distinctive pre-percolation mode of cell competition. These actin-rich protrusions were induced at heterotypic interfaces and could be generated by both DWO and MWO cells (**Figs. 5A,B**), suggesting that protrusion formation is a general response to cell-type mismatch. However, elimination events were strongly enriched for DWO-derived EpiArms during MWO removal, whereas most DWO removal events occurred independently of MWO-derived EpiArms (**Fig. 5D**). This asymmetry suggests that although both cell types can form protrusions, only DWO-derived EpiArms efficiently function as elimination machinery in this system. Their three-dimensional organization further supports a mechanical role: EpiArms extended beneath MWO cells, penetrated the basal region and locally deformed MWO cell nuclei, with nuclear shrinkage occurring preferentially during protrusion retraction (**Figs. 5C,C’,E**). These behaviours resemble epithelial arm-like structures used for cooperative apoptotic-cell clearance in zebrafish embryos^56^, and also recall filopodia-like protrusions that probe tissue mechanics during epithelial integration^57^. EpiArms may therefore combine recognition, mechanical probing, and force-generating functions at heterotypic interfaces. How these protrusions detect cellular mismatch, and why DWO-derived but not MWO-derived EpiArms are effective in elimination, remain important questions.

After percolation, MWO cell elimination occurred through a distinct lamellipodial squeezing mode. DWO cells surrounding MWO cells extended basal lamellipodia beneath the target cell, progressively closed the basal gap and subsequently constricted the mid-apical interface to extrude the MWO cell. This sequence is consistent with earlier work showing that neighbour-cell extending lamellipodia and contractile responses contribute to epithelial sealing and extrusion^18,20,58–61^. In our system, ERK activation in surrounding DWO cells preceded MWO cell rounding and accompanied extrusion, while acute ERK activation promoted lamellipodial extension at heterotypic interfaces (**Figs. 6F–J**). These observations support a model in which restored ERK spatio-temporal patterns after percolation help coordinate collective squeezing during MWO removal^34^. However, we note that the ERK dynamics observed here differ from previously described apoptosis-induced ERK waves that promote epithelial survival and sealing after cell death^24,28^. In those contexts, ERK activation is often discussed as a response to dying or extruding cells that protects neighbouring cells and restores barrier continuity. In the present system, ERK activity is embedded in a longer self-organized transition: DNA-damaged cells first suppress ERK patterning by disrupting mechanical connectivity, and ERK waves re-emerge only after cell competition restores a connected normal-cell network. Thus, ERK is not merely a downstream response to individual extrusion events, but part of a tissue-scale feedback loop linking connectivity, mechanochemical communication and elimination dynamics. This may explain why post-percolation extrusion is associated with ERK activation in neighbouring DWO cells, whereas pre-percolation elimination proceeds through protrusive EpiArm-mediated interactions under an intermediate, non-oscillatory ERK state.

More broadly, our findings reveal a form of feedback between local cellular interactions and global tissue topology. In classical percolation, local occupation or connectivity rules are typically fixed, and the global transition emerges when these local rules generate a spanning cluster^46,47,62^. In the epithelial system studied here, local interactions between normal and DNA-damaged cells drive the percolation transition by eliminating mechanically disruptive cells. Once the normal-cell network percolates, however, the resulting global topology changes the local rules of interaction by switching the dominant mode of cell elimination. This reciprocal loop, i.e., local competition drives global percolation and global percolation reshapes local competition, highlights a defining feature of living self-organized materials. Biological tissues can modify not only their connectivity, but also the behaviours that operate within a given connectivity state.

Several questions remain open. First, although ERK activity is a central readout and regulator of epithelial mechanochemical coordination, the transition described here is unlikely to be governed by ERK alone. Other pathways controlling actin polymerization, adhesion turnover, cortical contractility, cell–substrate coupling and DNA-damage responses are likely to contribute to both EpiArm dynamics and post-percolation squeezing. Second, the mechanical basis of EpiArm-mediated elimination remains to be studied. Direct measurements of protrusive force, target-cell deformation, adhesion disruption and local traction would clarify whether EpiArms primarily probe mechanical fitness, transmit forces, displace basal adhesions or initiate aberrant cell extrusion. Third, the percolation threshold identified here should be interpreted as an operational threshold in a finite experimental field of view rather than a universal constant; it will likely depend on cell type, tissue geometry, imaging scale, mixing ratio and the mechanical contrast between competing populations. Developing a comprehensive theoretical description of mechanochemical dynamics in such heterogeneous populations could further illuminate the underlying connections between physical properties and cell competition^63^. Finally, whether topology-dependent switching operates in vivo remains an important question. The murine epidermis, in which DNA-damage-associated cellular stress contributes to stem-cell competition and tissue ageing^64^ and ERK activity is dynamically patterned under physiological conditions^26^, offers a promising context in which to explore how competitive elimination may reshape epithelial connectivity and mechanochemical organization in living tissues.

## MATERIALS AND METHODS

### Cell lines and culture

Madin-Darby canine kidney (MDCK) cells were obtained from the RIKEN BioResource Center (RCB0995). MDCK cells stably expressing the nuclear-localized ERK FRET biosensor EKAREV-NLS (MDCK-ERK) and Lifeact-mCherry (MDCK-Lifeact-mCherry) were established as described previously^31,65^. Cells were maintained in Dulbecco’s modified Eagle’s medium (DMEM; Thermo Fisher Scientific, #11965092) supplemented with 10% heat-inactivated fetal bovine serum (FBS; Thermo Fisher Scientific, #A5670801) and 1% penicillin-streptomycin (10,000 U ml-1; Thermo Fisher Scientific, #15140148), hereafter referred to as complete medium. Cultures were maintained at 37°C in a humidified incubator with 5% CO2. Cells were routinely passaged before reaching overconfluence and were used for experiments after confirming normal epithelial morphology.

### Generation of DWO and MWO cells and co-culture assay

DMSO-washout (DWO) and mitomycin C-washout (MWO) cells were generated based on a previously described protocol for DNA-damage-induced epithelial cell competition^35^. Briefly, MDCK cells were trypsinized, collected by centrifugation and resuspended in complete medium. Cells were treated in suspension with either 8 µg ml^-1^ mitomycin C (MMC; Sigma-Aldrich, #M4287-2MG) or the corresponding volume of dimethyl sulfoxide (DMSO; Thermo Fisher Scientific) for 1 h at 37°C in a humidified 5% CO_2_ incubator. After treatment, cells were washed twice with fresh complete medium to remove residual MMC or DMSO. The time of reseeding after washout was defined as Day 0. For co-culture experiments, two cell populations were mixed immediately after washout at the indicated DWO:MWO ratios, including 50:1, 10:1 and 1:1, and seeded onto collagen-coated glass-bottom dishes. Unless otherwise stated, cells were seeded onto 35-mm dishes at 4×10^5^ cells per dish. To distinguish populations, MDCK-Lifeact-mCherry cells were prepared as either DWO or MWO populations, while MDCK-ERK cells served as the counterpart population. For long-term co-culture assays, cells were seeded in 24-well plates, and the medium was replaced daily with either fresh complete medium or drug-containing medium, as indicated.

### Antibodies, dyes, small molecules and recombinant proteins

The following primary antibodies (1:200 dilution), secondary antibodies (1:1000 dilution), and fluorescent dyes were used for immunofluorescence staining: rabbit monoclonal anti-cleaved caspase-3 (Cell Signaling Technology, #9664), rat monoclonal anti-E-cadherin (Thermo Fisher Scientific, #13-1900), rabbit polyclonal anti-gamma H2A.X (Abcam, #ab11174), rabbit polyclonal anti-phosphorylated myosin light chain (Abcam, #ab2480), mouse monoclonal anti-phosphorylated p53 (Cell Signaling Technology, #9286), goat anti-mouse IgG (H+L) Alexa Fluor 647 (Abcam, #ab150115), goat anti-rabbit IgG (H+L) Alexa Fluor 546 (Thermo Fisher Scientific, #A11035), goat anti-rabbit IgG (H+L) Alexa Fluor 647 (Abcam, # ab150079), goat anti-rat IgG (H+L) Alexa Fluor 647 (Thermo Fisher Scientific, #A21247), Hoechst 33342 (Thermo Fisher Scientific, #62249) for nuclear staining, and phalloidin-far-red (Thermo Fisher Scientific, #A22287) for F-actin staining.

The following small molecules and recombinant proteins were used where indicated: latrunculin A (Cayman Chemical, #10010630), mitomycin C (Sigma-Aldrich, #M4287), PD0325901 (Sigma-Aldrich, #PZ0162), TPA (12-O-tetradecanoylphorbol-13-acetate) (Sigma-Aldrich, #P8139), recombinant EGF (Sigma-Aldrich, #E9644).

### Immunofluorescence staining and confocal imaging

Cells were cultured on glass-bottom dishes (Iwaki, #3971-035), fixed with 4% paraformaldehyde (PFA) for 15 min at room temperature (23°C), and subsequently washed twice with phosphate-buffered saline (PBS) for 5 min each. Fixed cells were permeabilized with 0.1% Triton X-100 in PBS (PBST) for 5 min and subsequently washed with PBS. Samples were then blocked with 10% normal goat serum (Cell Signaling Technology, #5425S) for 2 h at 37°C. Cells were incubated with primary antibodies diluted in PBS overnight at 4°C. After three washes with PBS, samples were incubated with appropriate secondary antibodies and/or fluorescent staining dyes for 2 h at 37°C or overnight at 4°C. Following staining, samples were washed twice with PBS and stored in PBS at 4°C until imaging. Fluorescence images were acquired using an AX R confocal microscope (Nikon) equipped with a CFI Plan Apochromat VC 60X water-immersion objective (NA=1.2; Nikon, #MRD07602). For apical and basal cytoskeletal analyses, z-stacks were acquired through the monolayer and optical planes corresponding to the apical junctional region and basal cell-substrate interface were selected for analysis.

### Microdevice fabrication for through-hole stencils

NOA73 through-hole stencils were fabricated using a soft-lithography-based replica-molding strategy adapted from established microfabrication approaches^66^. UV-curable Norland Optical Adhesive 73 (NOA 73; Norland Products, Inc.) was used as the stencil material. Silicon master wafers containing hexagonal arrays of microwells were prepared by standard microfabrication. The microwell diameters were 3200, 1600, 800, 400, 200 and 100 µm, with an inter-well spacing of 150 µm and a depth of 100 µm. To generate polydimethylsiloxane (PDMS) micropillar molds, SYLGARD silicone elastomer base and curing agent were mixed at a 10:1 mass ratio, poured over the silicon master to a thickness of at least 3 mm, degassed under vacuum until no visible bubbles remained, and cured at 80°C for 1.5 h. The cured PDMS replica, containing micropillar arrays corresponding to the microwells of the silicon master, was gently peeled from the wafer and cut into pieces containing the desired pattern sizes.

To fabricate NOA73 stencils, the PDMS micropillar mold was placed on a flat PDMS base with the micropillars facing down and gently pressed to ensure close contact between the pillar tips and the base surface. NOA73 was introduced into the gap by capillary flow, and the mold– NOA73–base assembly was exposed to UV light at 20% power for 2 min. Close contact between the pillar tips and the PDMS base was maintained to prevent formation of residual NOA73 membranes across the through-holes. After curing, the NOA73 stencil was carefully peeled from the PDMS mold using fine forceps and cut to the desired size. Stencils were inspected before use to confirm complete through-hole opening, absence of residual membranes or tearing, and overall pattern integrity. Only intact stencils with uniform contact to the culture substrate were used for micropatterned co-culture assays.

### Stencil-based micropatterned co-culture assay

For micropatterning experiments, 27-mm glass-bottom dishes (Iwaki, #3970-035) were coated with type I collagen (Sigma-Aldrich, #C4243) diluted in 20 mM acetic acid to 30 µg ml^-1^ for 2 h and washed twice with PBS. Cured NOA73 stencils were attached to the collagen-coated glass-bottom dishes and sterilized under UV light in a biosafety cabinet for 10–15 min. Before cell seeding, complete medium was added to the stencil-mounted dishes, which were then degassed under vacuum at approximately 5 mmHg for 5 min to remove trapped air from the through-holes and improve uniform cell loading. To generate DWO cell islands, MDCK-ERK cells were seeded onto dishes fitted with NOA73 stencils at approximately 375 cells mm^-2^ and incubated for 16-24 h until confluency was reached within the stencil openings. The NOA73 stencil was then carefully and quickly removed using fine forceps to minimize detachment of the preformed DWO islands. MWO-treated MDCK-Lifeact-mCherry cells were subsequently seeded around the DWO islands at approximately 187 cells mm^-2^. After 2 h of incubation, dishes were gently shaken and aspirated to remove unbound cells and minimize nonspecific attachment of surrounding cells onto the preformed islands. Fresh complete medium was added, and co-cultures were maintained before live imaging.

### Live cell fluorescence imaging

Live cell fluorescence imaging was performed in FluoroBrite DMEM (Thermo Fisher Scientific, #1896701) supplemented with 10% FBS. Cells were maintained at 37°C in a humidified atmosphere with 5% CO₂ using a stage-top incubator (Tokai Hit) throughout imaging. Fluorescence images were acquired using an AX R confocal microscope (Nikon). For live ERK FRET imaging and large-field-of-view imaging, either a CFI Plan Fluor DLL 20X air objective lens (NA=0.5; Nikon, #MRH10205) or a CFI Plan Apochromat VC 20X air objective lens (NA=0.75; Nikon, #MRD70200) was used. For high-resolution 3D imaging, a CFI Plan Apochromat VC 60× water-immersion objective lens (NA 1.20; Nikon, #MRD07602) was used. For ERK FRET imaging, the donor fluorophore was excited with a 445 nm laser. Emission signals were collected using band-pass filters of 464–499 nm for the CFP donor channel and 518–551 nm for the FRET acceptor channel. For mCherry imaging, fluorescence was excited with a 561 nm laser. When CFP–YFP FRET and mCherry were imaged simultaneously, a custom multiband dichroic mirror was used to enable efficient detection of CFP, YFP, and mCherry emission signals. The dichroic mirror was selected to transmit fluorescence in the approximate emission ranges of 455–500 nm, 525–555 nm, and 580–630 nm, while minimizing transmission in the intervening excitation/reflection bands. Images were acquired using galvano scanning with two-line averaging. Long-term ERK FRET imaging was typically performed at 5- or 10-min intervals. Images were generally acquired at 512 × 512 pixels for standard live imaging and at 2048 × 2048 pixels for high-resolution imaging of Lifeact–mCherry-labelled actin protrusions.

### Image segmentation and cell tracking

Nuclear segmentation and single-cell tracking were performed using the TrackMate–Cellpose plugin^67^ implemented in ImageJ/Fiji (National Institutes of Health). For preprocessing, the FRET-channel images acquired from live imaging of MDCK-ERK or MDCK-Lifeact-mCherry cells were used as input. Images were first denoised using a 3 × 3 median filter to reduce shot noise. Pixel intensities were then normalized using contrast-limited adaptive histogram equalization (CLAHE) with default ImageJ parameters to enhance nuclear contrast for the object recognition. Nuclei were detected using the Cellpose detector within TrackMate. The target object diameter was set in the range of 15–20 µm for DWO cells and 25-30 µm for MWO cells, depending on the sample and imaging conditions. Detected nuclei were subsequently linked over time using TrackMate’s built-in tracking algorithms with default linking and gap-closing parameters. The resulting trajectories were used for downstream quantitative analyses.

### Particle image velocimetry

Flow fields were estimated from time-lapse images of nuclear CFP signals using particle image velocimetry (PIV) implemented in OpenPIV^68^. PIV was performed using an interrogation window of 32 pixels × 32 pixels, corresponding to 55.7 µm × 55.7 µm, with an overlap of 24 pixels. Displacements were calculated between image pairs separated by six frames, corresponding to a time lag of 60 min. MWO domains were identified by binarising the Lifeact–mCherry signal. This mask was used to define the spatial distribution of MWO cells, whereas PIV analysis itself was performed on the nuclear image data from the full co-culture field.

### Mean squared displacement analysis

MSD was calculated to quantify the persistence and diffusive properties of single-cell trajectories. Cell positions were obtained from time-lapse imaging and tracked over time to generate trajectories ***r***(*t*). The MSD was defined as:

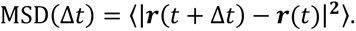

The angle brackets denote averaging over all cells and time points. MSD curves were computed over a range of time lags and plotted on a log–log scale. To characterize the mode of cell motion, MSD curves were fitted to a power-law relation: MSD(Δ*t*)∼(Δ*t*)*^α^*, where *α* is the scaling exponent. *α* = 1 indicates diffusive motion, *α* > 1 indicates super-diffusive (persistent) motion, and *α* < 1 indicates subdiffusive behaviour. The exponent *α* was estimated by linear regression of the log– log MSD curves over specified time intervals.

### Visualization and quantification of ERK activity

For visualization and quantification of ERK activity, raw CFP and FRET images were first denoised using a 3 x 3 median filter. Background fluorescence was subtracted independently from the CFP and FRET channels at each time point, and FRET/CFP ratio images were generated using custom MATLAB scripts (MathWorks). For pseudocolour visualization, hue represents the FRET/CFP ratio, whereas brightness corresponds to the FRET channel intensity. Single-cell ERK activity was quantified using cell positions obtained from Cellpose-based segmentation and TrackMate-based tracking, as described above. The centroid coordinates of each segmented object were exported from TrackMate and imported into custom MATLAB scripts. For each cell, a 5 x 5-pixel region of interest centered on the centroid was defined, and the median FRET/CFP ratio within this region was calculated. This median FRET/CFP ratio was used as the ERK activity value for each individual cell at each time point. The median value was used to reduce the influence of local noise and outlier pixels within the measurement region.

### Image-based counting of labelled cells in co-culture

The number of Lifeact-mCherry-labelled cells in co-culture was quantified from fluorescence images using a Cellpose–TrackMate-based segmentation workflow, as described above. Prior to segmentation, labelled-cell nuclear images were extracted using a custom MATLAB script. Briefly, images were processed by median filtering, contrast-limited adaptive histogram equalization, Otsu thresholding, and morphological closing to generate a binary mask for the labelled population. The mask was then used to isolate the corresponding labelled-cell signal and generate processed images for downstream segmentation.

Processed images were segmented using Cellpose through TrackMate. The Cellpose nuclear model was used with an expected object diameter of 15 µm for Day 1.5 and Day 3.5 images. For Day 5.5 and Day 7.5 images, the expected diameter was set to 10 µm, as larger diameter settings underestimated cell numbers. For low-intensity images, including Day 7.5 datasets, CLAHE was applied twice before segmentation to improve nuclear detection. Segmented objects were exported as TrackMate spots, and the number of detected spots was used as the number of labelled cells. The total cell number in each field of view was obtained from the corresponding original nuclear images using the same Cellpose–TrackMate workflow. Images used for automated quantification covered fields of approximately 2500 µm × 2500 µm. Manual cell counts were also performed in randomly selected 400 µm × 400 µm windows from the original images and compared with the automated segmentation results to validate the automated counting. The resulting labelled-cell counts and total cell counts were used to calculate the fraction of Lifeact-mCherry-labelled cells in each co-culture condition.

### Estimation of percolation probability

Percolation probability was quantified from time-lapse images of DWO–MWO co-cultures using a custom MATLAB workflow. For each time point, images were first converted into binary masks representing DWO-occupied regions. Images were processed by median filtering, Otsu thresholding, morphological opening, and morphological closing, followed by mask inversion when required to define the DWO-positive domain. To evaluate local network connectivity, each field of view was subdivided into overlapping square windows. The window size was set to half of the full image width, corresponding to approximately 1.63 mm × 1.63 mm images. Windows were shifted with a step size corresponding to 20% of the window size. For each window, the DWO occupancy fraction, *ϕ*, was calculated as the fraction of pixels assigned to the DWO-positive domain. Percolation was assessed for each window by connected-component analysis using 4-connectivity. A window was scored as percolated when at least one connected DWO-positive component spanned the window from left to right or from top to bottom. Each window was therefore assigned a binary percolation score, with 1 indicating percolated and 0 indicating non-percolated. To obtain the percolation probability *P*(*ϕ*), all windows from all time points were pooled and binned according to their DWO occupancy fraction using 0.02-wide ϕ intervals. For each bin, *P*(*ϕ*) was calculated as the mean of the binary percolation scores within that bin. Percolation curves were generated independently for each dataset, smoothed using a three-point moving average, and then averaged across datasets. When required, the critical occupancy fraction was operationally defined as the value of *ϕ* at which *P*(*ϕ*) reached 0.95 and was estimated by interpolation. This analysis estimates the probability that DWO-positive regions form a locally spanning network at a given occupancy fraction, providing an operational measure of percolation-like connectivity in finite experimental fields of view.

### Measurement of protrusion frequency at cell interfaces

Actin-rich protrusions at homotypic and heterotypic interfaces were quantified from live imaging of co-cultures containing Lifeact–mCherry-labelled and unlabelled cells. Interfaces were identified based on the boundary between the two cell populations visualized by the Lifeact–mCherry signal. Protrusions were defined as Lifeact–mCherry-positive extensions projecting toward or into the unlabelled cell population and clearly distinguishable from local cortical fluctuations. Only protrusive structures longer than 20 µm were included in the analysis. For each field of view, the number of interface cells was counted at the initial time point. Protrusion formation was then assessed over a 180-min imaging window, and the total number of protrusion events appearing during this period was manually counted. Protrusions that persisted across consecutive frames were counted as a single event. The protrusion frequency was calculated by dividing the total number of protrusion events by the number of interface cells in the same field of view.

### Classification of EpiArm-associated cell elimination

Cell elimination events were identified from time-lapse phase-contrast images based on the appearance of rounded cell morphology during removal. For each elimination event, the projected area of the rounded/removing cell was defined from the phase-contrast image at the time of removal. The corresponding mCherry channel was then inspected to determine whether EpiArms from heterotypic cells extended into the projected area of the removing cell. Events were scored as EpiArm-positive when mCherry-labelled EpiArms overlapped with the projected area of the removing cell, and EpiArm-negative otherwise. The position of each removing cell was also classified according to whether it was located at a heterotypic DWO–MWO interface/edge. Based on these two binary classifications, each event was assigned to one of four categories: EpiArm+/Edge+, EpiArm+/Edge−, EpiArm−/Edge+, or EpiArm−/Edge−. The proportion of each category was then calculated from the total number of elimination events analyzed.

### Mathematical modeling and simulations

To model the ERK dynamics we simulate the dynamics of a vertex model with mechanochemical coupling to ERK^41^. Briefly, the vertex model energy per cell is given as

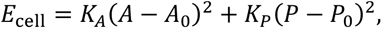

with area *A*, target area *A*_0_, area elasticity *K_A_*, perimeter *P*, target perimeter *P*_0_, and perimeter elasticity *K_P_*. The target area of one cell is coupled to its ERK level ℰ with coupling strength *α* and temporal delay *τ_A_* via a cell-specific ordinary differential equation,

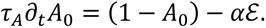

The cell-specific ERK offset from its reference value is coupled to the strain, i.e., the area, via

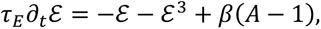

with the ERK-sensitivity *β* and delay time scale *τ_E_*. We use the parameters from the previous paper^41^, which have been fit to describe ERK waves in MDCK monolayers, except that we neglect random cell movements to concentrate on the network’s propensity to become unstable to waves. To incorporate the different cellular motion patterns, we assume as target index of DWO cells *p*_0_ = *P*_0_/ C*A*_0_ = 3.9, i.e., they are fluid, while for MWO cells we assume *p*_0_ = 3.8, corresponding to mechanically solid-like cells. A mechanochemical coupling strength of (*αβ*) = 2(*αβ*)*_crit_* was found to yield experimentally realistic critical domain sizes, where (*αβ*)*_crit_* is the critical coupling of this system^41^. For the MWO cells, we assumed *α* = *β* = 0, i.e., they are mechanochemically inactive, but the wave attenuation behavior was verified to also hold for partially mechanically inactive MWO cells, i.e., (*αβ*) = 0.5(*αβ*)*_crit_* (**Fig. S3**). The monolayers were generated as random Voronoi tessellations, with cell centres determined by random sequential adsorption. For simulations in circular domains, boundary nodes were fixed on the circle, whereas periodic boundary conditions were used for all other simulations. For percolation-transition simulations, the MWO network pattern was extracted from the MWO cell mask by binarisation and used to select corresponding cells in a periodic random Voronoi tessellation. Areas were non-dimensionalised by the estimated mean DWO cell area, *A*_ref_ = 400 µm^2^. Cell velocities were calculated from the displacement of cell centres of mass. Time was non-dimensionalised using the ERK timescale adopted previously^41^, i.e., *τ_E_* = 6 min. Because this fixed timescale was retained for rescaling, the dimensionalised oscillation period and cell speeds show a slight mismatch with the experimental values.

### Statistical analysis

The number of cells analyzed (n) and the number of biological replicates (N) are reported in the figure legends and/or Table S1. No statistical methods were used to predetermine sample sizes, and no randomization was performed. Statistical tests, sample sizes and corresponding p values are described in the figure legends and/or **Table S1**. Unless otherwise stated, two-tailed tests were used and p < 0.05 was considered statistically significant. Significance levels are denoted as follows: *p < 0.05, **p < 0.01, ***p < 0.001 and n.s., not significant.

### Software and data visualization

Digital image processing, quantitative analysis and visualization were performed using R2024b (MathWorks), ImageJ/Fiji (National Institutes of Health). Truncated violin plots were generated using a MATLAB function from MATLAB File Exchange^69^.

## ACKNOWLEDGEMENTS

This work was supported by Singapore Ministry of Education (MOE) Academic Research Fund (AcRF) Tier 2 (MOE-T2EP30223–0010), the National Research Foundation, Singapore (NRF) under its Mid-sized Grant (NRF-MSG-2023–0001), and the Mechanobiology Institute (MBI) at the National University of Singapore funded through the National Research Foundation, Singapore and the Ministry of Education, Singapore under the Research Centre of Excellence programme to T.H. This work also received funding from the ERC under the European Union’s Horizon 2020 research and innovation programme (No. 851288) to E.H. Microfabrication of through-hole stencils and image-processing computation were supported by MBI’s Microfabrication Core and Computational Core Facility, respectively. We thank all members of the Grant-in-Aid for Transformative Research Areas (A) ‘Multicellular Autonomy’ for fruitful discussions, and members of the Hirashima and Hannezo labs for their valuable comments on the manuscript.

## DATA AVAILABILITY

Data collected for this study will be made publicly available via the digital institutional repository of the National University of Singapore.

## CODE AVAILABILITY

Relevant code for this study is available at the following GitHub repository: https://github.com/oliverdrozdowski/HeterogeneousERK

## AUTHOR CONTRIBUTIONS

Conceptualization: A.N., T.H.

Methodology: A.N., O.M.D., E.V., E.H., T.H.

Investigation: A.N., O.M.D., E.V., S.W., T.P.N., T.H.

Data curation: A.N., O.M.D., E.V., T.H.

Formal analysis: A.N., O.M.D., T.H.

Writing – original draft: A.N., O.M.D., E.V., E.H., T.H. Writing – review & editing: E.H., T.H.

Supervision: E.H., T.H. Funding acquisition: E.H., T.H. Resources: E.H., T.H.

Visualization: A.N., O.M.D., E.H., T.H.

## SUPPLEMENTAL FIGURES

**Figure S1.**
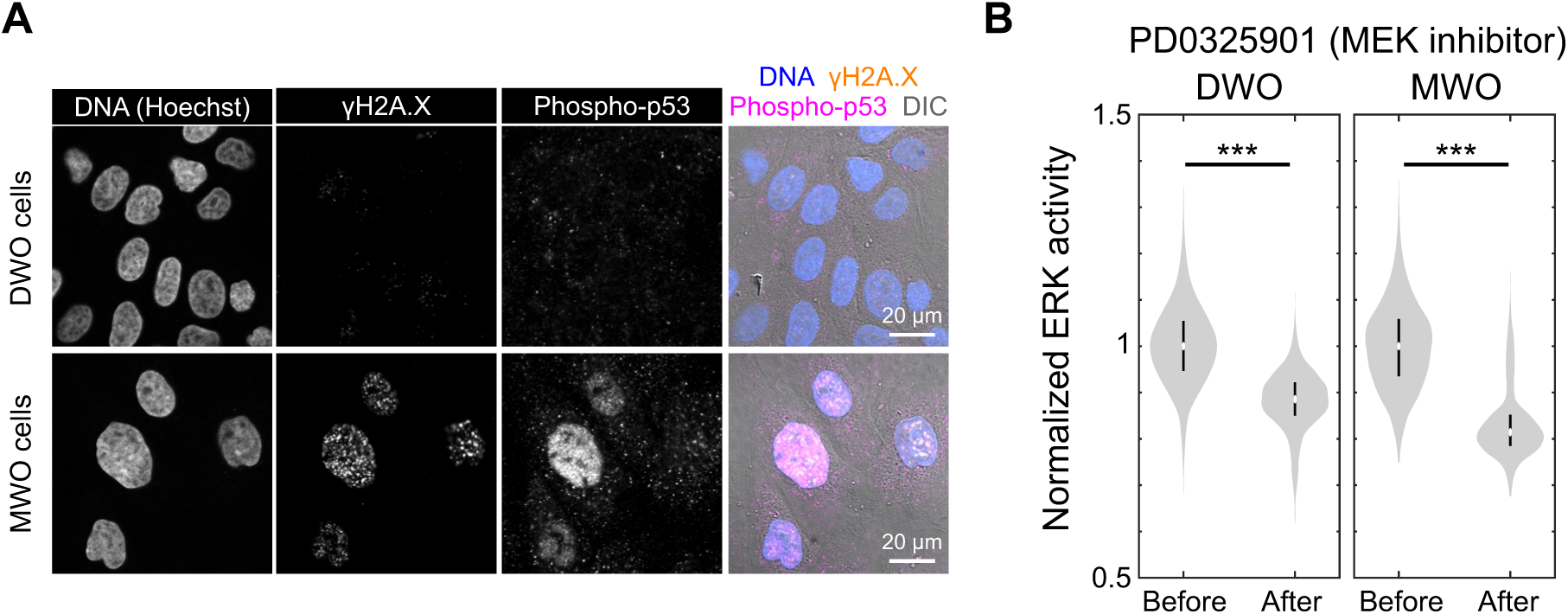
Characterization of DWO and MWO cells, related to Figure 1. (A) Immunostaining for γ-H2A.X and phosphorylated p53, with Hoechst nuclear counterstaining, in DWO (top) and MWO (bottom) cells on Day 1. Scale bars, 20 µm. (B) ERK activity before and after treatment with 1 µM PD0325901, a MEK inhibitor, in DWO (left) and MWO (right) cells. “Before” values were pooled from five time points before treatment (−25, −20, −15, −10, and −5 min; DWO: n = 4,934; MWO: n = 2,512; N = 3 each). “After” values were pooled from two time points after treatment (80 and 85 min; DWO: n = 1,767; MWO: n = 760; N = 3 each). Two-sample t-test, p < 1e-32 for both conditions.

**Figure S2.**
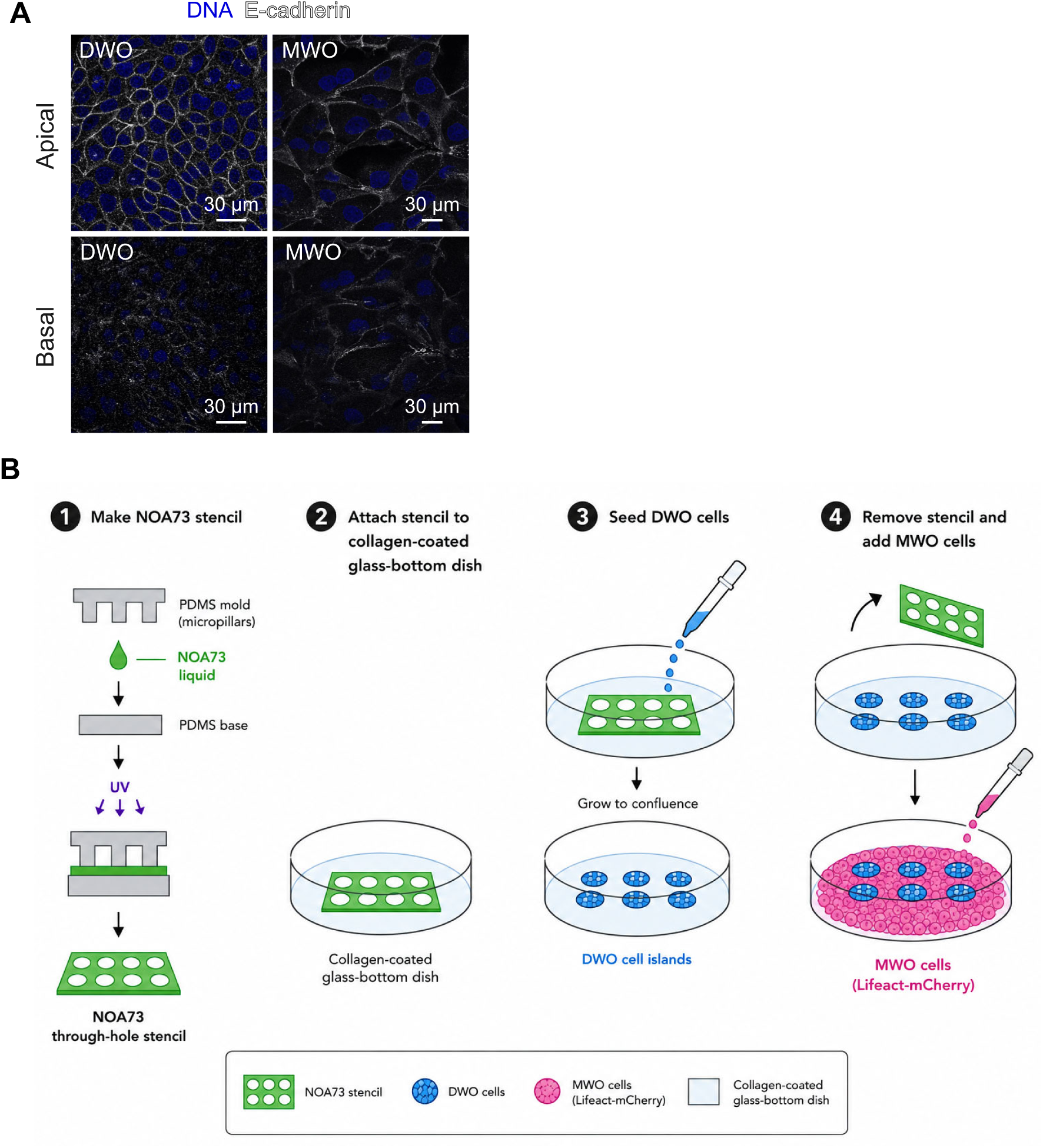
Mechanical characterization and micro-stencil assay, related to Figure 2. (A) Apical (top) and basal (bottom) distribution of E-cadherin in DWO (left) and MWO (right) cells, with Hoechst nuclear staining. Scale bars, 30 µm. (B) Microdevice fabrication for through-hole stencils and stencil-based co-culture assay. Briefly, NOA73 was molded using a PDMS micropillar mold and UV-cured to generate through-hole stencils. The stencil was attached to a collagen-coated glass-bottom dish, and DWO cells were seeded into the through-holes to form spatially defined cell islands. After stencil removal, MWO cells were seeded into the surrounding area to generate DWO cell islands embedded within an MWO cell population. See Materials and Method sections: ‘Microdevice fabrication for through-hole stencils’ and ‘Stencil-based micropatterned co-culture assay’ for the details.

**Figure S3.**
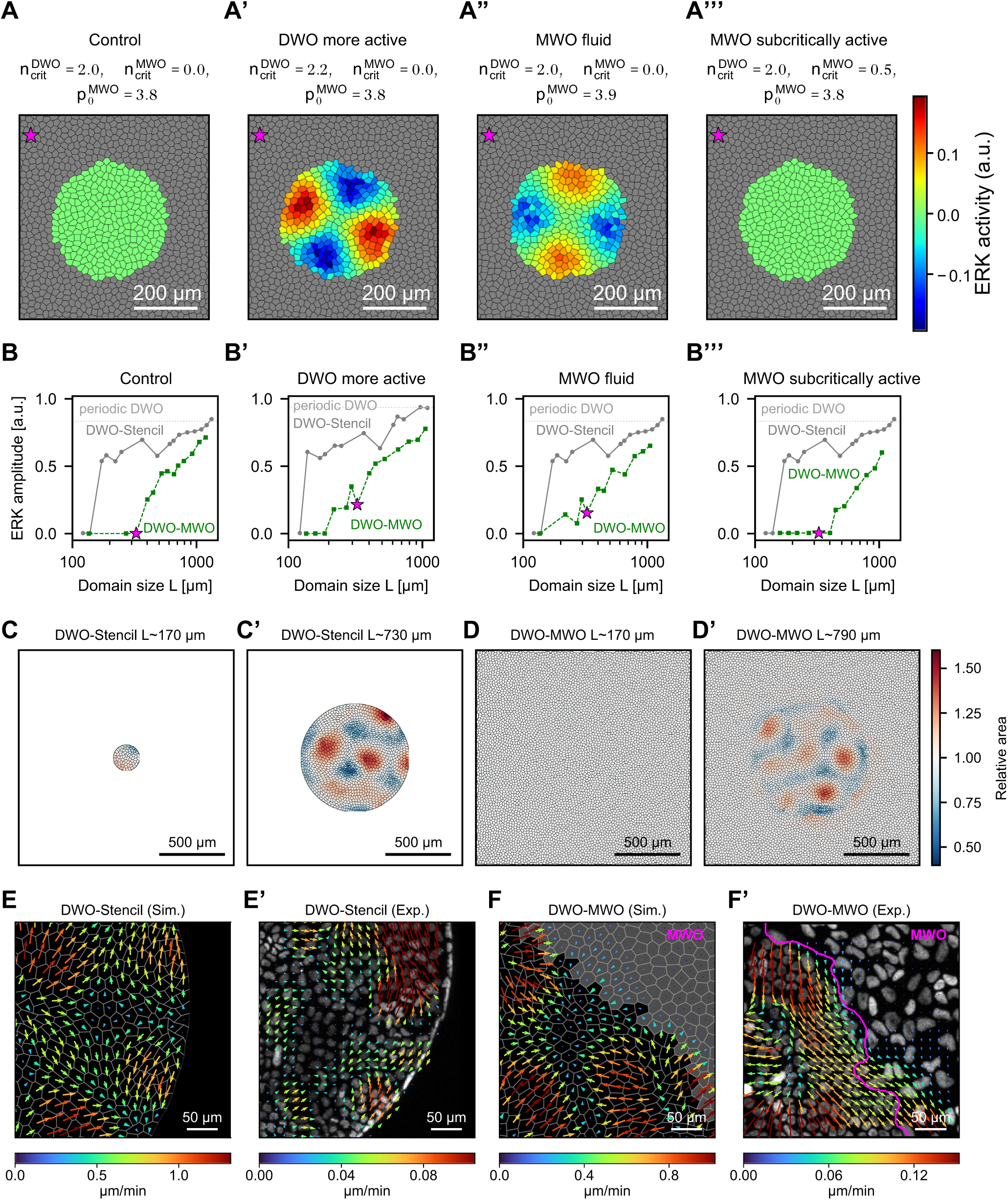
Sensitivity and boundary effects of wave attenuation in the theoretical vertex model, related to Figure 2. (A–A’’’) ERK activity pattern of DWO cells surrounded by MWO cells (grey) for parameters used in Fig. 2N (A), with increased mechanochemical coupling strength n_crit_ quantifying the relative size of mechanochemical coupling parameter *αβ* compared to the critical threshold (see methods) (A’), and for an increased tissue fluidity of the MWO cells characterized through the target shape index *p^M^*^W0^ = *P*_0_/C*A*_0_, characterized by target cell perimeter P_0_ and target area A_0_ (A’’), and for mechanochemically subcritical MWO cells (A’’’). Scale bars, 200 µm. (B–B’’’) The ERK amplitude curves for the DWO-Stencil, the DWO-MWO and the periodic DWO case as in Fig. 2N, corresponding to the cases in (A–A’’’). (C,C’) The relative areas (without ERK coupling areas are 1.0) of cells in the DWO-Stencil case for small (C) and large (C’) patterns. Scale bars, 500 µm. (D,D’) The relative areas of cells in the DWO-MWO case for small (D) and large (D’) islands of DWO cells surrounded by MWO cells. Scale bars, 500 µm. (E,E’) Simulated velocity fields of DWO cells on the stencil (E) and corresponding flow fields obtained via particle image velocimetry (PIV) (E’). Scale bars, 50 µm. (F,F’) Simulated velocity fields of DWO cells surrounded by MWO cells, with MWO cells shown in grey (F), and corresponding flow fields obtained via PIV, with the domain boundary highlighted in magenta (F’). In the flow fields, relative arrow sizes and arrow colours indicate image-specific relative flow velocities. Scale bars, 50 µm.

**Figure S4.**
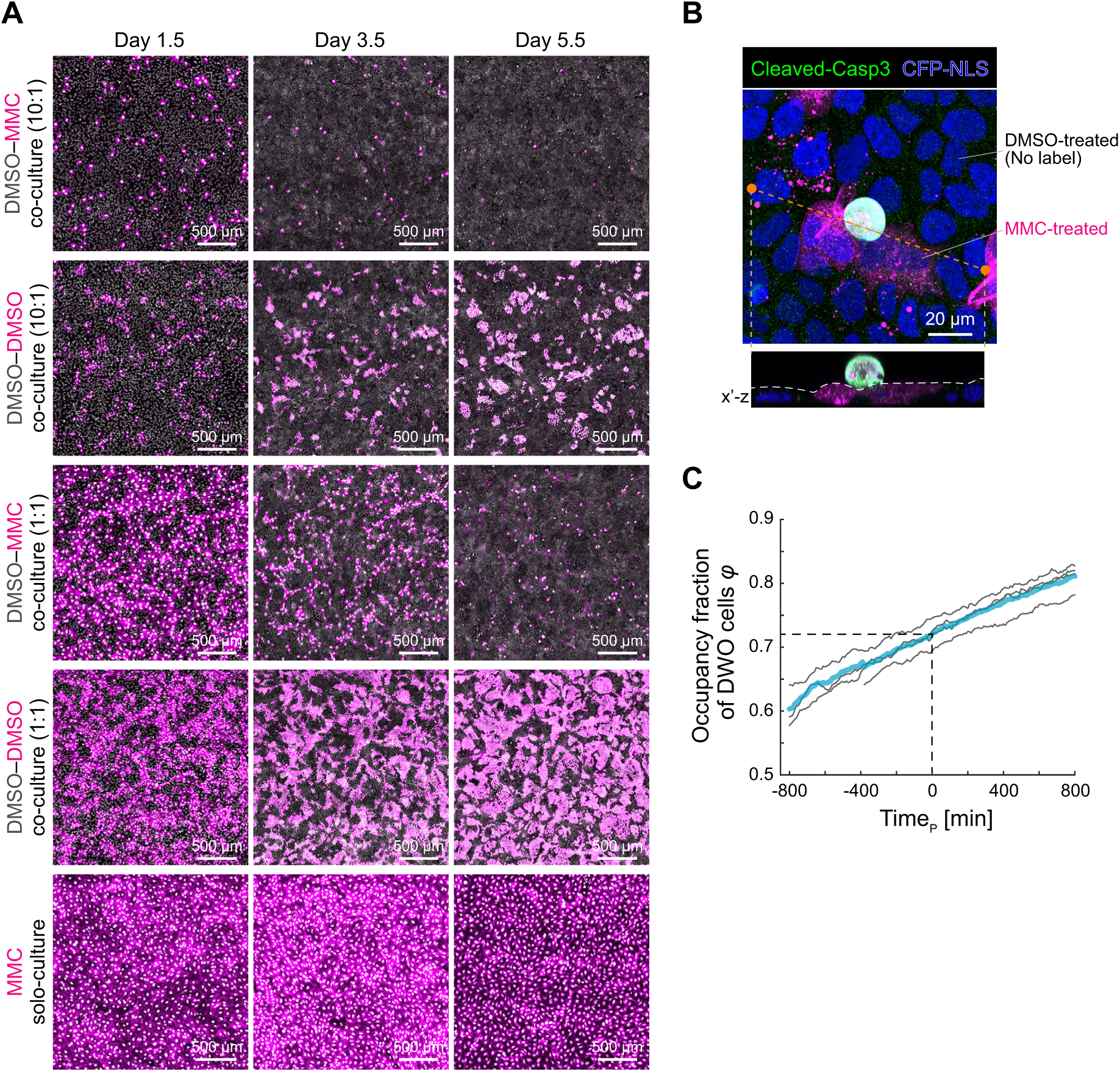
Characterization for competitive elimination of MWO cells, related to Figure 3. (A) Long-term co-culture assay under different culture conditions: 10:1 co-culture (first and second rows), 1:1 co-culture (third and fourth rows), and MWO solo culture (fifth row). Cell populations labelled with Lifeact–mCherry are shown in magenta and indicated on the left. Images in the third and fourth rows are the same as those shown in Fig. 3A and are included here for comparison. Scale bars, 500 µm. (B) Immunostaining for cleaved caspase-3 (green), with CFP nuclear counterstaining (blue) in DWO–MWO co-culture. MWO cells are labelled with Lifeact–mCherry (magenta). The orange dashed line indicates the x’ axis used for orthogonal reconstruction; the lower x’–z view shows the apico-basal plane. Scale bar, 20 µm. (C) Occupancy fraction of DWO cells (φ) as a function of time aligned to the critical percolation point. Time 0 denotes the critical percolation time (Time_p_=0), at which the percolation probability reaches *P* = 0.95. Gray lines represent individual samples, and the light blue line indicates the mean. N=4.

**Figure S5.**
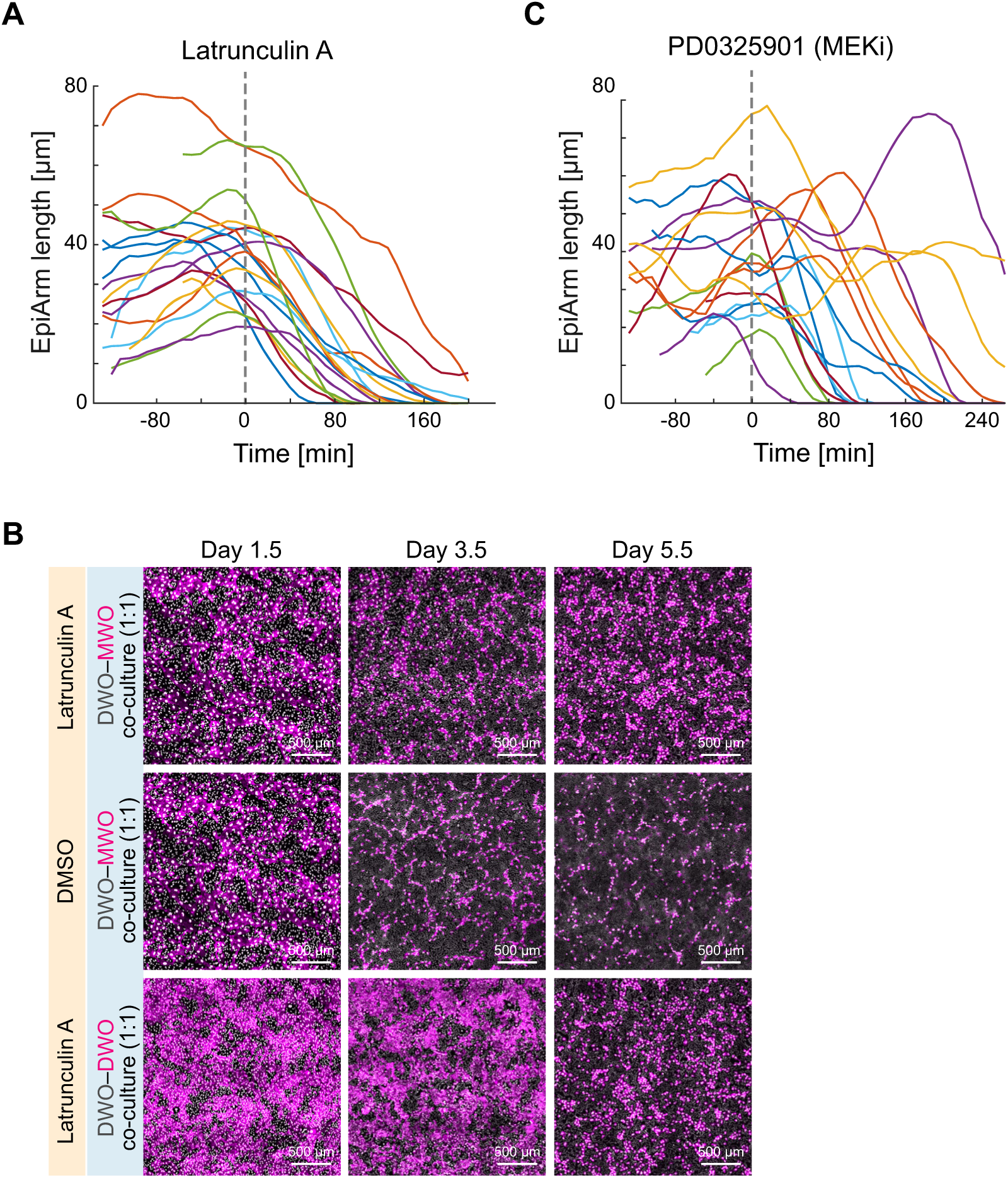
Pharmacological perturbation of EpiArm dynamics and DWO–MWO cell competition, related to Figure 5. (A) EpiArm length in DWO cells over time before and after treatment with 100 nM Latrunculin A in DWO–MWO co-culture. Time 0 denotes the time of treatment. n=19 from N=3. (B) Long-term 1:1 DWO–MWO co-culture assay with 100 nM Latrunculin A treatment (first and third rows) or DMSO control (second row). Cell populations labelled with Lifeact–mCherry are shown in magenta and indicated on the left. Drug- or vehicle-containing medium was refreshed daily. Scale bars, 500 µm. (C) EpiArm length in DWO cells over time before and after treatment with 1 µM PD0325901, a MEK inhibitor, in DWO–MWO co-culture. Time 0 denotes the time of treatment. n=18 from N=3.

**Figure S6.**
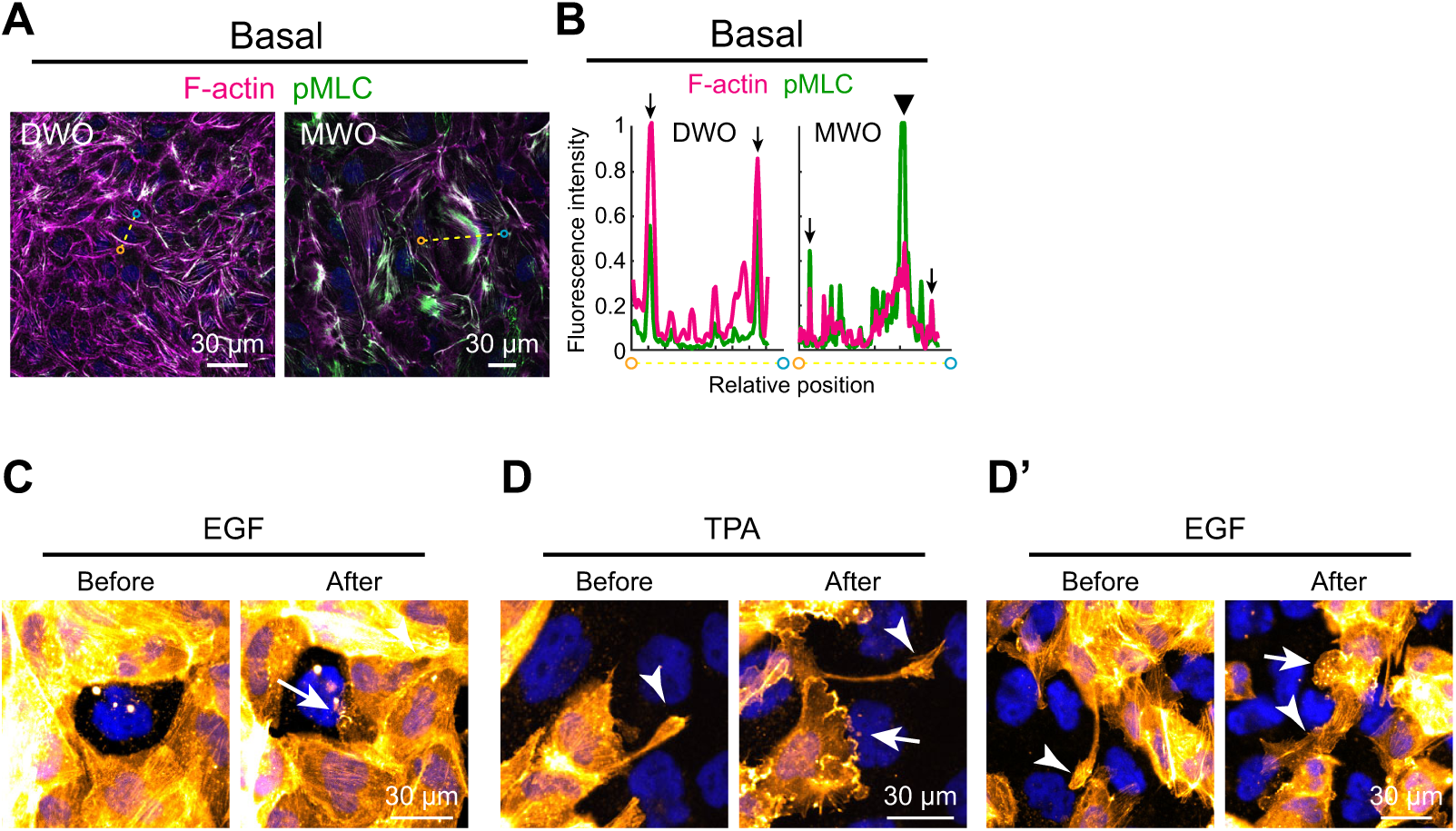
ERK activation induces lamellipodial extension, related to Figure 6. (A,B) Basal distribution of F-actin (magenta) and pMLC (green) in DWO (left) and MWO (right) cells during pre-percolation. (A) Fluorescence images. (B) Corresponding intensity profiles along the yellow dotted line indicated in (A). Arrowheads indicate the cell cortex; black triangles denote pronounced accumulation of F-actin and pMLC beneath the nucleus. In contrast to the apical depletion, MWO cells exhibited marked basal accumulation of F-actin and pMLC beneath the nucleus and along the basal membrane, whereas DWO cells maintained the typical junctional enrichment of these contractile components. Scale bars, 30 µm. (C) Snapshots of F-actin organization before and after treatment with 100 ng mL^-1^ EGF during the post-percolation phase. DWO cells are labelled with Lifeact–mCherry (orange) and surround an unlabelled MWO cell. Nuclei are counterstained with CFP (blue). The arrow indicates lamellipodial extension. Scale bar, 30 µm. (D,D’) Snapshots of F-actin organization before and after treatment with 10 nM TPA (D) or 100 ng mL^-1^ EGF (D’) during the pre-percolation phase. DWO cells are labelled with Lifeact–mCherry (orange), whereas MWO cells are unlabelled. Nuclei are counterstained with CFP (blue). Arrows indicate lamellipodial extensions, and arrowheads indicate EpiArms. Scale bars, 30 µm.

## SUPPLEMENTAL VIDEOS

**Video 1. ERK activity dynamics in DWO solo culture and DWO–MWO co-cultures.**

Time-lapse images of ERK activity on Day 2 in DWO solo culture and DWO–MWO co-cultures at ratios of 50:1, 10:1, and 1:1, shown from left to right. MWO cells are shown in magenta. ERK activity is colour-coded as in **Figure 1C**.

**Video 2. ERK activity dynamics in MWO solo culture.**

Time-lapse images of ERK activity in MWO solo culture on Day 2. ERK activity is colour-coded as in **Figure 1C**.

**Video 3. ERK activity dynamics in small confined DWO clusters surrounded by MWO cells**

Time-lapse images of ERK activity in confined DWO clusters with a typical domain size of L ≈ 220 µm, surrounded by MWO cells shown in magenta. ERK activity is colour-coded as in **Figure 2I**.

**Video 4. ERK activity dynamics in large confined DWO clusters surrounded by MWO cells**

Time-lapse images of ERK activity in confined DWO clusters with a typical domain size of L ≈ 600 µm, surrounded by MWO cells shown in magenta. ERK activity is colour-coded as in **Figure 2I’**.

**Video 5. ERK and area dynamics of simulated DWO-Stencil and DWO-MWO experiments**

Simulated time-lapse of ERK levels (top) and relative areas (bottom) in DWO cells confined within the circular stencils (DWO-Stencil) or surrounded by MWO cells under periodic boundary conditions (DWO–MWO). Corresponds to **Figure 2M**.

**Video 6. ERK activity dynamics during percolation in DWO–MWO co-culture.**

Time-lapse images of ERK activity in 1:1 DWO–MWO co-culture on Day 2.5. MWO cells are shown in magenta. Time 0 is aligned to the critical percolation point. ERK activity is colour-coded as in **Figure 4A**.

**Video 7. Simulated ERK dynamics in pre- and post-percolation tissue patterns.**

Simulated time-lapse of spatially distributed ERK levels (top) and single-cell ERK levels (bottom) in DWO–MWO mixtures during the pre- and post-percolation phases. The analyzed single cells are marked by magenta stars in the top panels. Gray cells denote MWO cells. The DWO–MWO spatial arrangements were extracted from the time-lapse data shown in **Figure 4A** and were held fixed during the simulations. Corresponds to **Figures 4C,D**.

**Video 8. MWO cell elimination driven by DWO-derived EpiArms.**

Time-lapse imaging of MWO cell elimination driven by DWO-derived EpiArms. Left: phase-contrast images. Right: corresponding fluorescence images showing DWO cells labelled with Lifeact–mCherry in orange and unlabelled MWO cells with nuclei counterstained in blue. Time 0 is defined as the onset of nuclear shrinkage preceding extrusion. Corresponds to **Figure 5E**.

**Video 9. EpiArm retraction after Latrunculin A treatment.**

Time-lapse images showing morphological changes in DWO EpiArms (orange) before and after treatment with 100 nM Latrunculin A. MWO cells are unlabelled, with nuclei counterstained in blue. Time 0 denotes the time of treatment. Corresponds to **Figures 5F,G**.

**Video 10. EpiArm dynamics after MEK inhibition.**

Time-lapse images showing morphological changes in DWO EpiArms (orange) before and after treatment with 1 µM PD0325901, a MEK inhibitor. MWO cells are unlabelled, with nuclei counterstained in blue. Time 0 denotes the time of treatment. Corresponds to **Figures 5H,I**.

**Video 11. MWO cell elimination by surrounding cell squeezing during the post-percolation phase**

Time-lapse imaging of MWO cell elimination driven by squeezing from surrounding DWO cells during the post-percolation phase. DWO cells are labelled with Lifeact–mCherry (orange), whereas the central MWO cell is unlabelled. Nuclei are counterstained with CFP (blue). Left and middle panels show basal and middle-plane F-actin signals in DWO cells; right panels show phase-contrast images at the apical plane. Time 0 is defined as the onset of MWO nuclear shrinkage. Corresponds to **Figure 6C**.

**Video 12. ERK activity dynamics during MWO cell elimination in the post-percolation phase**

Time-lapse imaging showing elimination of an MWO cell (magenta) and ERK activity in surrounding DWO cells during the post-percolation phase. Time 0 is defined as the onset of MWO cell shrinkage. ERK activity is colour-coded as in **Figure 6F**.

**Video 13. Lamellipodium extension induced by TPA treatment.**

Time-lapse images showing morphological changes in DWO EpiArms (orange) before and after treatment with 10 nM TPA. MWO cells are unlabelled, with nuclei counterstained in blue. Time 0 denotes the time of treatment. Corresponds to **Figure S6D**.

**Video 14. Lamellipodium extension induced by EGF treatment.**

Time-lapse images showing morphological changes in DWO EpiArms (orange) before and after treatment with 100 ng mL^-1^ EGF. MWO cells are unlabelled, with nuclei counterstained in blue. Time 0 denotes the time of treatment. Corresponds to **Figure S6D′**.

## REFERENCES

1. Bailles, A., Gehrels, E. W. & Lecuit, T. Mechanochemical Principles of Spatial and Temporal Patterns in Cells and Tissues. Annu. Rev. Cell Dev. Biol. 38, 321–347 (2022).

2. Brückner, D. B. & Hannezo, E. Tissue Active Matter: Integrating Mechanics and Signaling into Dynamical Models. Cold Spring Harb Perspect Biol 17, a041653 (2025).

3. Hannezo, E. & Heisenberg, C.-P. Mechanochemical Feedback Loops in Development and Disease. Cell 178, 12–25 (2019).

4. Lenne, P.-F. et al. Roadmap for the multiscale coupling of biochemical and mechanical signals during development. Phys. Biol. 18, 041501 (2021).

5. Alert, R. & Trepat, X. Physical Models of Collective Cell Migration. Annu. Rev. Condens. Matter Phys. 11, 77–101 (2020).

6. Campàs, O., Noordstra, I. & Yap, A. S. Adherens junctions as molecular regulators of emergent tissue mechanics. Nat Rev Mol Cell Biol 25, 252–269 (2024).

7. Ladoux, B. & Mège, R.-M. Mechanobiology of collective cell behaviours. Nat Rev Mol Cell Biol 18, 743–757 (2017).

8. Xi, W., Saw, T. B., Delacour, D., Lim, C. T. & Ladoux, B. Material approaches to active tissue mechanics. Nat Rev Mater 4, 23–44 (2018).

9. Balasubramaniam, L. et al. Dynamic forces shape the survival fate of eliminated cells. Nat. Phys. 21, 269–278 (2025).

10. Clavería, C. & Torres, M. Cell Competition: Mechanisms and Physiological Roles. Annu. Rev. Cell Dev. Biol. 32, 411–439 (2016).

11. Dwivedi, V. K. et al. Replication stress promotes cell elimination by extrusion. Nature 593, 591–596 (2021).

12. Maruyama, T. & Fujita, Y. Cell competition in vertebrates — a key machinery for tissue homeostasis. Current Opinion in Genetics & Development 72, 15–21 (2022).

13. Ohsawa, S., Vaughen, J. & Igaki, T. Cell Extrusion: A Stress-Responsive Force for Good or Evil in Epithelial Homeostasis. Developmental Cell 44, 284–296 (2018).

14. Van Neerven, S. M. & Vermeulen, L. Cell competition in development, homeostasis and cancer. Nat Rev Mol Cell Biol 24, 221–236 (2023).

15. Wagstaff, L. et al. Mechanical cell competition kills cells via induction of lethal p53 levels. Nat Commun 7, 11373 (2016).

16. Aoki, K. et al. Mechano-gradients drive morphogen-noise correction to ensure robust patterning. Sci. Adv. 10, eadp2357 (2024).

17. Drozdowski, O. M., Kocameşe-Tamgacι, B., Boonekamp, K. E., Boutros, M. & Schwarz, U. S. Cell Bulging and Extrusion in a Three-Dimensional Bubbly Vertex Model for Curved Epithelial Sheets. Phys. Rev. X 16, 021023 (2026).

18. Kocgozlu, L. et al. Epithelial Cell Packing Induces Distinct Modes of Cell Extrusions. Current Biology 26, 2942–2950 (2016).

19. Krueger, D. et al. Epithelial tension controls intestinal cell extrusion. Science 389, eadr8753 (2025).

20. Le, A. P. et al. Adhesion-mediated heterogeneous actin organization governs apoptotic cell extrusion. Nat Commun 12, 397 (2021).

21. Pothapragada, S. P., Gupta, P., Mukherjee, S. & Das, T. Matrix mechanics regulates epithelial defence against cancer by tuning dynamic localization of filamin. Nat Commun 13, 218 (2022).

22. Schoenit, A., Mège, R.-M. & Ladoux, B. Biophysical principles of cell competition and elimination. Trends in Cell Biology S096289242600067X (2026) doi:10.1016/j.tcb.2026.04.009.

23. De Simone, A. et al. Control of osteoblast regeneration by a train of Erk activity waves. Nature 590, 129–133 (2021).

24. Gagliardi, P. A. et al. Collective ERK/Akt activity waves orchestrate epithelial homeostasis by driving apoptosis-induced survival. Developmental Cell 56, 1712–1726.e6 (2021).

25. Hirashima, T. & Matsuda, M. ERK-mediated curvature feedback regulates branching morphogenesis in lung epithelial tissue. Current Biology 34, 683–696.e6 (2024).

26. Hiratsuka, T. et al. Intercellular propagation of extracellular signal-regulated kinase activation revealed by in vivo imaging of mouse skin. eLife 4, e05178 (2015).

27. Ishii, M., Tateya, T., Matsuda, M. & Hirashima, T. Retrograde ERK activation waves drive base-to-apex multicellular flow in murine cochlear duct morphogenesis. eLife 10, e61092 (2021).

28. Valon, L. et al. Robustness of epithelial sealing is an emerging property of local ERK feedback driven by cell elimination. Developmental Cell 56, 1700–1711.e8 (2021).

29. Aoki, K. et al. Propagating Wave of ERK Activation Orients Collective Cell Migration. Developmental Cell 43, 305–317.e5 (2017).

30. Boocock, D., Hino, N., Ruzickova, N., Hirashima, T. & Hannezo, E. Theory of mechanochemical patterning and optimal migration in cell monolayers. Nat. Phys. 17, 267– 274 (2021).

31. Hino, N. et al. ERK-Mediated Mechanochemical Waves Direct Collective Cell Polarization. Developmental Cell 53, 646–660.e8 (2020).

32. Crozet, F. & Levayer, R. Emerging roles and mechanisms of ERK pathway mechanosensing. Cell. Mol. Life Sci. 80, 1–19 (2023).

33. Hirashima, T., Hino, N., Aoki, K. & Matsuda, M. Stretching the limits of extracellular signal-related kinase (ERK) signaling — Cell mechanosensing to ERK activation. Current Opinion in Cell Biology 84, 102217 (2023).

34. Nayak, A. N. & Hirashima, T. Tug-of-war via ERK signaling pathway for tissue organization – ERK activation to force generation. Current Opinion in Cell Biology 85, 102249 (2023).

35. Kozyrska, K. et al. p53 directs leader cell behavior, migration, and clearance during epithelial repair. Science 375, eabl8876 (2022).

36. Tomasz, M. Mitomycin C: small, fast and deadly (but very selective). Chemistry & Biology 2, 575–579 (1995).

37. Harvey, C. D., et al. A genetically encoded fluorescent sensor of ERK activity. Proc. Natl. Acad. Sci. U.S.A. 105, 19264–19269 (2008).

38. Komatsu, N. et al. Development of an optimized backbone of FRET biosensors for kinases and GTPases. MBoC 22, 4647–4656 (2011).

39. Villars, A. & Levayer, R. Collective effects in epithelial cell death and cell extrusion. Current Opinion in Genetics & Development 72, 8–14 (2022).

40. Cook, A. W. & Toseland, C. P. The roles of nuclear myosin in the DNA damage response. The Journal of Biochemistry 169, 265–271 (2021).

41. Boocock, D., Hirashima, T. & Hannezo, E. Interplay between Mechanochemical Patterning and Glassy Dynamics in Cellular Monolayers. PRX Life 1, 013001 (2023).

42. Muthukrishnan, S. et al. Glassy dynamics in active epithelia emerge from an interplay of mechanochemical feedback and crowding. Nat Commun 17, 7391 (2026).

43. Yu, P., Zhang, R. & Li, B. Cell Stiffness-Mediated Mechanochemical Waves in Three-Dimensional Tissues. Phys. Rev. Lett. 135, 108401 (2025).

44. Yu, P., Li, Y., Fang, W., Feng, X.-Q. & Li, B. Mechanochemical dynamics of collective cells and hierarchical topological defects in multicellular lumens. Sci. Adv. 10, eadn0172 (2024).

45. Baker, N. E. Emerging mechanisms of cell competition. Nat Rev Genet 21, 683–697 (2020).

46. Li, M. et al. Percolation on complex networks: Theory and application. Physics Reports 907, 1–68 (2021).

47. Saberi, A. A. Recent advances in percolation theory and its applications. Physics Reports 578, 1–32 (2015).

48. Lenne, P.-F. & Trivedi, V. Sculpting tissues by phase transitions. Nat Commun 13, 664 (2022).

49. Thukral, S., Dey, B. & Wang, Y. Integrating Tissue and Cytoplasmic Rigidity Transitions During Morphogenesis. Development, Growth & Differentiation 67, 378–394 (2025).

50. Mongera, A. et al. A fluid-to-solid jamming transition underlies vertebrate body axis elongation. Nature 561, 401–405 (2018).

51. Park, J.-A. et al. Unjamming and cell shape in the asthmatic airway epithelium. Nature Mater 14, 1040–1048 (2015).

52. Pinheiro, D. & Mitchel, J. Pulling the strings on solid-to-liquid phase transitions in cell collectives. Current Opinion in Cell Biology 86, 102310 (2024).

53. Bi, D., Lopez, J. H., Schwarz, J. M. & Manning, M. L. A density-independent rigidity transition in biological tissues. Nature Phys 11, 1074–1079 (2015).

54. Petridou, N. I., Corominas-Murtra, B., Heisenberg, C.-P. & Hannezo, E. Rigidity percolation uncovers a structural basis for embryonic tissue phase transitions. Cell 184, 1914–1928.e19 (2021).

55. Rustarazo-Calvo, L. et al. Adhesion-driven rigidity transition decoupled from density-driven jamming triggers epithelial organization in embryonic tissues. Nat. Phys. 10.1038/s41567-026-03276-6 (2026) doi:10.1038/s41567-026-03276-6.

56. Hoijman, E. et al. Cooperative epithelial phagocytosis enables error correction in the early embryo. Nature 590, 618–623 (2021).

57. Ventura, G. et al. Multiciliated cells use filopodia to probe tissue mechanics during epithelial integration in vivo. Nat Commun 13, 6423 (2022).

58. Bonfim-Melo, A. et al. Rapid lamellipodial responses by neighbor cells drive epithelial sealing in response to pyroptotic cell death. Cell Reports 38, 110316 (2022).

59. Kira, A. et al. Apoptotic extracellular vesicle formation via local phosphatidylserine exposure drives efficient cell extrusion. Developmental Cell S1534580723002411 (2023) doi:10.1016/j.devcel.2023.05.008.

60. Otani, T., Nguyen, T. P., Kinoshita, N., Fujimori, T. & Furuse, M. Competitive elimination of ZO-1/ZO-2-deficient cells regulates epithelial barrier homeostasis. Preprint at 10.1101/2025.01.10.632304 (2025).

61. Takeuchi, Y. et al. Calcium Wave Promotes Cell Extrusion. Current Biology 30, 670–681.e6 (2020).

62. Sahimi, M. Applications of Percolation Theory. vol. 213 (Springer International Publishing, Cham, 2023).

63. Li, X., Das, A. & Bi, D. Mechanical Heterogeneity in Tissues Promotes Rigidity and Controls Cellular Invasion. Phys. Rev. Lett. 123, 058101 (2019).

64. Liu, N. et al. Stem cell competition orchestrates skin homeostasis and ageing. Nature 568, 344–350 (2019).

65. Wp, S. et al. Transient contractility attenuation reprograms epithelial cells into a protrusion-driven state that drives tissue fluidization. Preprint at 10.64898/2026.03.23.713577 (2026).

66. Xia, Y. & Whitesides, G. M. Soft Lithography. Angewandte Chemie International Edition 37, 550–575 (1998).

67. Stringer, C. & Pachitariu, M. Cellpose3: one-click image restoration for improved cellular segmentation. Nat Methods 22, 592–599 (2025).

68. Liberzon, A., et al. OpenPIV/openpiv-python: fixed windows conda-forge failure with encoding. Zenodo 10.5281/ZENODO.3566451 (2019).

69. Bechtold, B., Fletcher, P., Seamusholden & Gorur-Shandilya, S. bastibe/Violinplot-Matlab: A Good Starting Point. Zenodo 10.5281/ZENODO.4559847 (2021).

